# Co-expression network analysis reveals repression of *Igf2bp2* by REV-ERBβ in skeletal muscle

**DOI:** 10.64898/2026.05.13.724827

**Authors:** Vishnu A.M., Qing Zhang, Shriyansh Srivastava, Kevin B. Koronowski, Ashutosh Srivastava

## Abstract

The circadian clock genes *Bmal1* and *Nr1d1/2* (REV-ERBα/β) regulate skeletal muscle metabolism and homeostasis, yet the precise genes and mechanisms involved remain incompletely understood. Here, we perform Weighted Gene Co-expression Network Analysis (WGCNA) on skeletal muscle circadian transcriptomes with varying *Bmal1* operational status to identify genes central to muscle circadian function. The largest WGCNA module, potentially under *Bmal1* regulation, contains clock and muscle-specific output genes governed hierarchically by hub genes including *Igf2bp2*, an RNA-binding protein involved in muscle progenitor growth and maintenance. *Igf2bp2* expression is rhythmic in mouse and human muscle and functional experiments in muscle-specific *Bmal1* knockout mice show that Igf2bp2 is upregulated by loss of *Bmal1* at ZT8 and negatively correlated with *Nr1d2*, suggesting de-repression through REV-ERBβ as a regulatory mechanism. Luciferase reporter experiments in cultured myotubes show that REV-ERBβ, but not REV-ERBα, represses *Igf2bp2* transcription and that repression is mediated by non-canonical GCC motifs in the *Igf2bp2* promoter region. Together, these findings uncover a circadian *Nr1d2-Igf2bp2* regulatory axis linking transcriptional and post-transcriptional regulation in skeletal muscle, with implications for muscle homeostasis.

**Highlights:**

- *Igf2bp2* clusters with *Nr1d2* (*Rev-erbβ*) in circadian co-expression network
- *Bmal1* or *Rev-erbɑ/β* knockout upregulates *Igf2bp2* in muscle
- *Igf2bp2* is rhythmic in WT muscle but arrhythmic in clock mutant muscle
- REV-ERBβ represses *Igf2bp2* transcription in myotubes
- REV-ERBβ repression requires GCC motifs in the *Igf2bp2* promoter

**Graphical Abstract:** 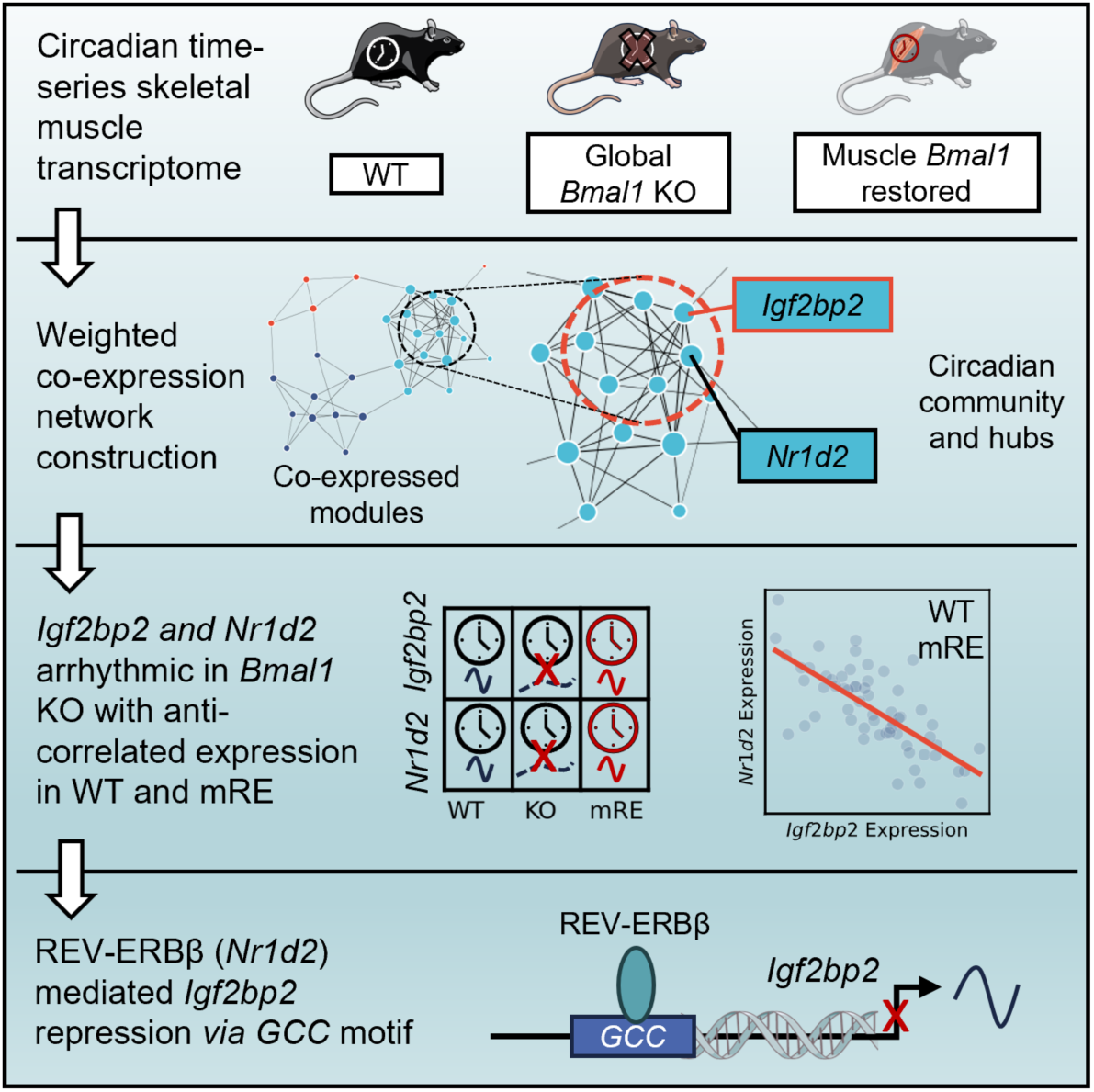

## Introduction

Circadian rhythms are endogenous biological oscillations with a period of approximately 24 hours that regulate diverse physiological processes in most living organisms (Patke et al., 2020). These ∼24h cycles are generated by intrinsic molecular clocks and are entrained to environmental cues (Stevenson et al., 2025). At the molecular level, circadian timekeeping relies on an autoregulatory transcriptional-translational feedback loop (TTFL) (Takahashi, 2017). In mammals, the positive limb of the TTFL consists of the transcription factors BMAL1 and CLOCK, which heterodimerize and drive transcription of core clock genes including Period (*Per1/2/3*) and Cryptochromes (*Cry1/2*). The PER and CRY proteins accumulate in the cytoplasm, translocate into the nucleus, and repress CLOCK:BMAL1 activity, thereby completing the negative feedback loop. Additionally, an auxiliary loop involving REV-ERBs (*Nr1d1/2*) and RORs further reinforces rhythmic oscillation by binding to the RORE motifs (Takahashi, 2017). Importantly, these core-clock proteins are central to a well-organized system of metabolic regulation and their disruption has been implicated in chronic metabolic disorders (Bass & Takahashi, 2010; Neves et al., 2022; Patke et al., 2020; Rijo-Ferreira & Takahashi, 2019; Takahashi, 2016, 2017; R. Zhang et al., 2014). Although the core circadian mechanism is conserved throughout the body, molecular clocks also mediate tissue-specific functions in a coherent manner (Kumar et al., 2024; Mure et al., 2018; Sica et al., 2026; Viggars et al., 2024; R. Zhang et al., 2014).

Since the circadian system is a multicomponent, highly interconnected network, a systems-level perspective is necessary to understand these complex regulatory mechanisms (Bechtel, 2024; Ueda, 2007). Importantly, publicly available high-throughput circadian transcriptomics datasets are key resources that provide an opportunity to understand the regulatory mechanisms at scale. A recent study used Weighted Gene Co-expression Network Analysis (WGCNA) to identify adaptive molecular mechanisms involved in calorie restriction at different time points on different days post calorie-restriction in liver and white adipose tissues (Pak et al., 2026). Despite their effectiveness in revealing biological insights, WGCNA studies on circadian transcriptomes are limited and have focused on the brain and liver, or global cycling genes (Bhargava et al., 2015; Brown et al., 2017; J. Li et al., 2022; Santos et al., 2026; Simak et al., 2019).

Of note, skeletal muscle has emerged as an important tissue to understand the crosstalk between circadian regulation and metabolism, primarily due to fundamental role of circadian rhythms in energy metabolism, glucose homeostasis, and exercise physiology (Dyar et al., 2013; Harfmann et al., 2016; Hodge et al., 2015; Perrin et al., 2018; Schroder et al., 2015). Interestingly, circadian transcriptomics studies of mouse skeletal muscle estimate that ∼4-16% of the skeletal muscle transcriptome is rhythmically expressed, reflecting both core clock activity and additional mechanisms that result in rhythmic transcription (Dyar et al., 2013; McCarthy et al., 2007). Muscle-specific clock-controlled genes include the myogenic regulator *Myod1* (Myogenic Differentiation 1) (involved in the myogenic lineage determination) (J. L. Andrews et al., 2010; Hodge et al., 2019; X. Zhang et al., 2012), key glycolytic enzymes such as *Hk2*, *Pfkm*, and *Pdk4*, *Pgc-1β* (Involved in regulation of lipid metabolism) (Harfmann et al., 2016; McCarthy et al., 2007; Smith et al., 2023), and the sarcomeric protein *Tcap* (Hodge et al., 2019; Riley et al., 2022). Disruption or loss of these genes results in the development of metabolic conditions such as impaired glucose tolerance, reduced insulin responsiveness, altered energy metabolism, and reduction in muscle mass, highlighting their importance in muscle physiology. Among the core clock genes, *Bmal1* is crucial for circadian regulation in skeletal muscle, and its loss has been shown to significantly disrupt the circadian rhythms and subsequent effects on muscle metabolism and functionality (Aoyama et al., 2021; Chaikin et al., 2025; Chatterjee et al., 2013; Dyar et al., 2013, 2018; Ehlen et al., 2017; Gao et al., 2020; Gutierrez-Monreal et al., 2024; Harfmann et al., 2016; Hodge et al., 2015; Schiaffino et al., 2016). Furthermore, other clock genes, *Clock*, *Per*, *Cry*, and *Nr1d1/2* also modulate time-dependent exercise effects, muscle maintenance, and metabolism *via* diverse mechanisms and pathways (J. L. Andrews et al., 2010; Y. Chen et al., 2026; Dyar et al., 2018; Hao et al., 2023; Jordan et al., 2017; Katoku-Kikyo et al., 2021; Kennaway et al., 2007; J. Liu et al., 2025; Mansingh et al., 2024).

While these studies have established key roles of circadian genes in skeletal muscle physiology, the complete map of transcriptional mechanisms governing this coordinated regulation is not clear. In light of this, methods like WGCNA (Langfelder & Horvath, 2008) provide an opportunity to decipher the unknown regulatory mechanisms.

A recent study investigated the interaction between liver and muscle clocks by generating mice which expressed *Bmal1* in liver, muscle (mRE), or both tissues, in an otherwise *Bmal1*-deficient animal (KO) (Smith et al., 2023), finding that liver and muscle clocks coordinate systemic glucose metabolism in tandem with daily feeding cycles. Here, leveraging transcriptomic datasets from muscle clock mutant mice, we constructed co-expression networks and performed WGCNA to identify the molecular mechanisms that underlie circadian metabolism in skeletal muscle. Using this framework, we found an interesting co-expression module containing regulatory hubs that displayed loss of rhythmicity in the KO genotype and gain of rhythms in mRE, suggesting a potential circadian regulation. With this analysis we report a novel mechanism of *Nr1d2* dependent regulation of a RNA-binding protein, *Igf2bp2* that was validated through qPCR and luciferase reporter assays. We propose that this *Nr1d2-Igf2bp2* axis could be crucial to the time-of-day dependent regulation of glucose metabolism, insulin-signaling, and RNA processing in skeletal muscles, and requires further experimental investigation.

## Results

### Co-expression network analysis identifies muscle gene modules specific to *Bmal1* status

We performed hierarchical clustering of gastrocnemius muscle circadian transcriptomes from wild-type (WT), *Bmal1* knockout (KO), and *Bmal1* muscle-reconstituted (mRE) mice (GSE197726) (Smith et al., 2023) to assess differences in global gene expression, and found a distinct grouping of samples according to genotype (Fig. 1A). While the WT and KO samples formed coherent clusters, mRE samples collected at ZT0, ZT4, and ZT20 tended to be more variable (Fig. 1A). Notably, the original study showed that muscle-specific *Bmal1* reconstitution only partially rescued rhythms, which could explain the variability among the mRE samples (Smith et al., 2023). Nevertheless, clustering indicated that the expression profiles of genotypes were different from each other. To further examine the sources of variation in the expression pattern, we performed principal component analysis (PCA) on the expression matrix across all samples. The first two principal components (PC1 and PC2) captured ∼35% of the total transcriptomic variance. Importantly, PC1 clearly separated KO samples from WT and mRE samples, indicating genotype as the dominant source of variation (Fig. 1B).

**Figure 1.**
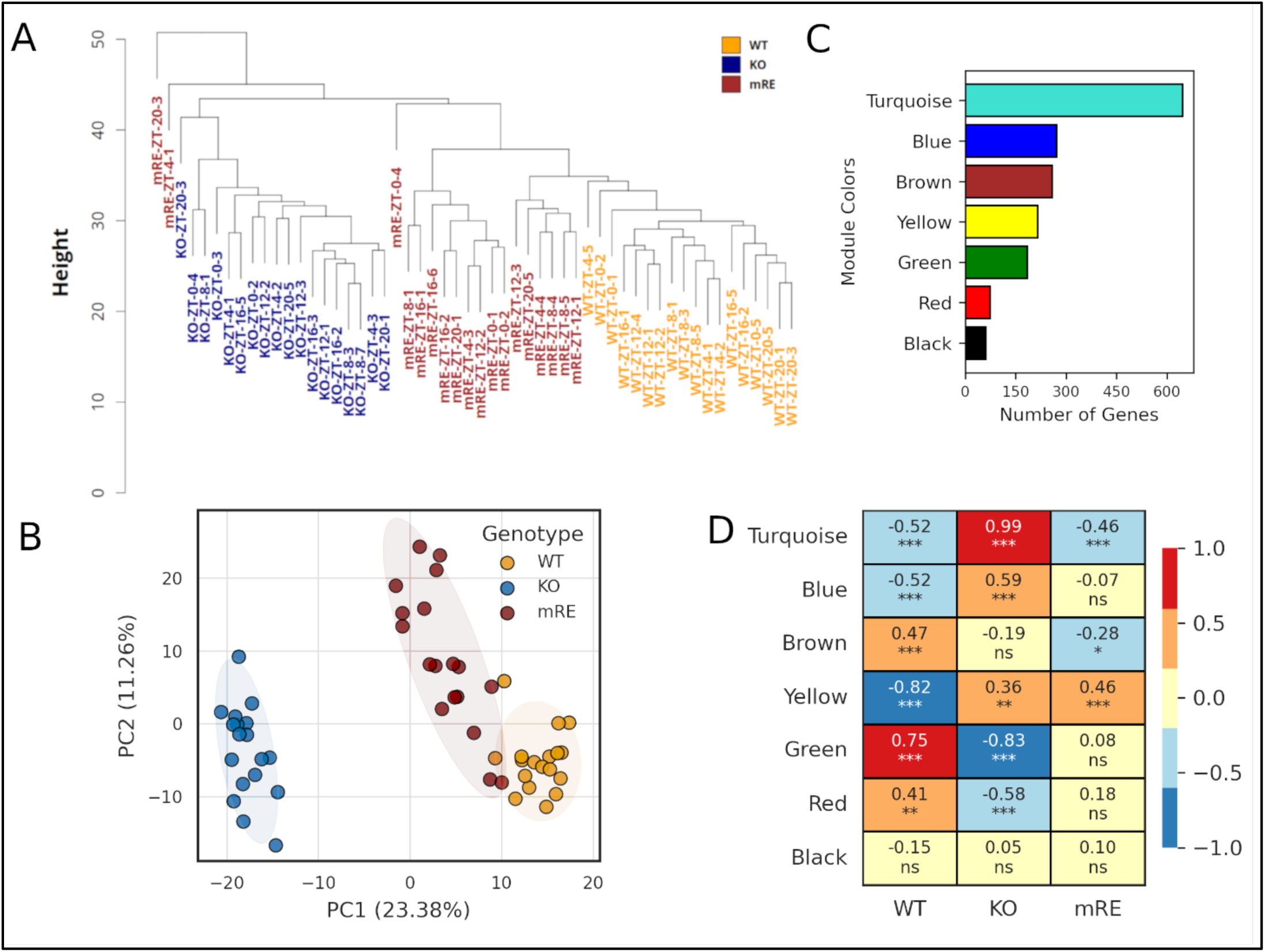
Global gene expression pattern and co-expression network reconstruction of circadian time-series bulk RNA-seq data. **(A)** Hierarchical clustering dendrogram of samples based on normalized gene expression profiles, showing segregation of WT, KO, and mRE groups (GSE197726). **(B)** Principal component analysis (PCA) of transcriptomic data demonstrating clear separation among WT, KO, and mRE samples along the first two principal components (PC1 and PC2). **(C)** Distribution of genes across WGCNA co-expression modules. Grey module has been removed for clarity. **(D)** Heatmap of module-trait correlations showing Pearson correlation coefficients between module eigengenes and different genotypes (WT, KO, mRE). Color scale represents correlation strength (red = positive, blue = negative), and asterisks indicate statistical significance (*p < 0.05, **p < 0.01, ***p < 0.001).

To understand the emerging gene co-expression patterns in the three genotypes, we performed WGCNA (see Methods), which identifies clusters of genes that have correlated expression and participate in the same biological pathway (Langfelder & Horvath, 2008). Clustering of this network revealed eight distinct modules. We focused on seven modules containing 1,715 genes that showed clear clustering based on co-expression (Fig. 1C). The remaining 12,203 genes could not be assigned to a proper co-expression module (grey module) and were excluded from further analysis. To understand the association of the co-expressed gene modules with genotypes, we calculated correlation between the expression of the module eigengene and genotype. Module eigengene represents overall expression of genes in a module. The module eigengenes for turquoise, blue, and yellow modules showed significant negative correlation with WT (p < 0.001), and positive correlation with KO (turquoise and blue: p < 0.001; yellow: p < 0.01) (Fig. 1D). In addition, the turquoise module eigengene was also negatively correlated with mRE (p < 0.001), whereas the blue module did not show a significant association. In contrast, the yellow module eigengene was positively correlated with mRE (p < 0.001). Importantly, the genotype specific correlation patterns distinguished functionally relevant module behavior. For instance, the turquoise module exhibited negative correlation with WT and mRE, but positive correlation with KO, suggesting that expression of genes in this module are restored towards the WT upon muscle-specific *Bmal1* reconstitution. In contrast, the yellow module remained negatively correlated with WT and positively correlated with both KO and mRE, indicating that these genes are not rescued by muscle-specific *Bmal1* reconstitution (Fig. 1D).

Conversely, the brown, green and red module eigengenes exhibited positive correlation with WT (brown and green, p < 0.001; red, p < 0.01) and a negative correlation with the KO genotype (green and red, p < 0.001; brown, n.s.). The green and red module eigengenes showed weak, non-significant positive correlations, while brown module eigengene exhibited negative correlation with the mRE genotype (p < 0.05), and the black module did not show any significant correlation across genotypes (Fig. 1D). Taken together, the module eigengenes revealed modules containing genes that were dysregulated in KO and (1) restored by muscle *Bmal1* (turquoise), (2) trending toward partial restoration (blue, green, red), or (3) not restored (yellow). To investigate muscle clock transcription further, we focused our analysis on the turquoise module, reasoning that it is likely to contain potentially novel clock-driven genes centrally involved in muscle function.

### Core clock genes cluster with muscle-specific regulatory hubs in the co-expression network

The significant co-expression exclusive to the turquoise module could be biologically meaningful since it contained the core-clock and well-established clock-controlled genes (see Supplementary Data 1). Next, we hypothesized that the turquoise module contained genes, established or potentially novel, that directly interact with the core circadian genes. To assess if there is another level of organization within this module, we performed community detection using the Louvain algorithm (Blondel et al., 2008). We identified eight communities. Inspection of community membership revealed that community 6 (hereafter referred to as ‘circadian community’) contained core clock genes such as *Clock*, *Per2*, *Per3*, *Cry1*, *Nr1d2*, and other known clock-controlled genes in the skeletal muscle (*Dbp, Tef, Nfil3, Tcap*), suggesting a highly coordinated expression pattern (see Supplementary Data 1). To locate the specific gene hubs that play a central role within the circadian community, we performed network connectivity, gene significance (GS), and module membership (MM) analyses across all communities (see Methods), identifying 28 hub genes (see Table S1). Of note, five of the 28 hubs were present in the circadian community including *Nr1d2*, *Per3*, *Tcap*, *Igf2bp2*, and *Gm40841* (Fig. 2A, also see Supplementary Data 1 and Table S1).

**Figure 2.**
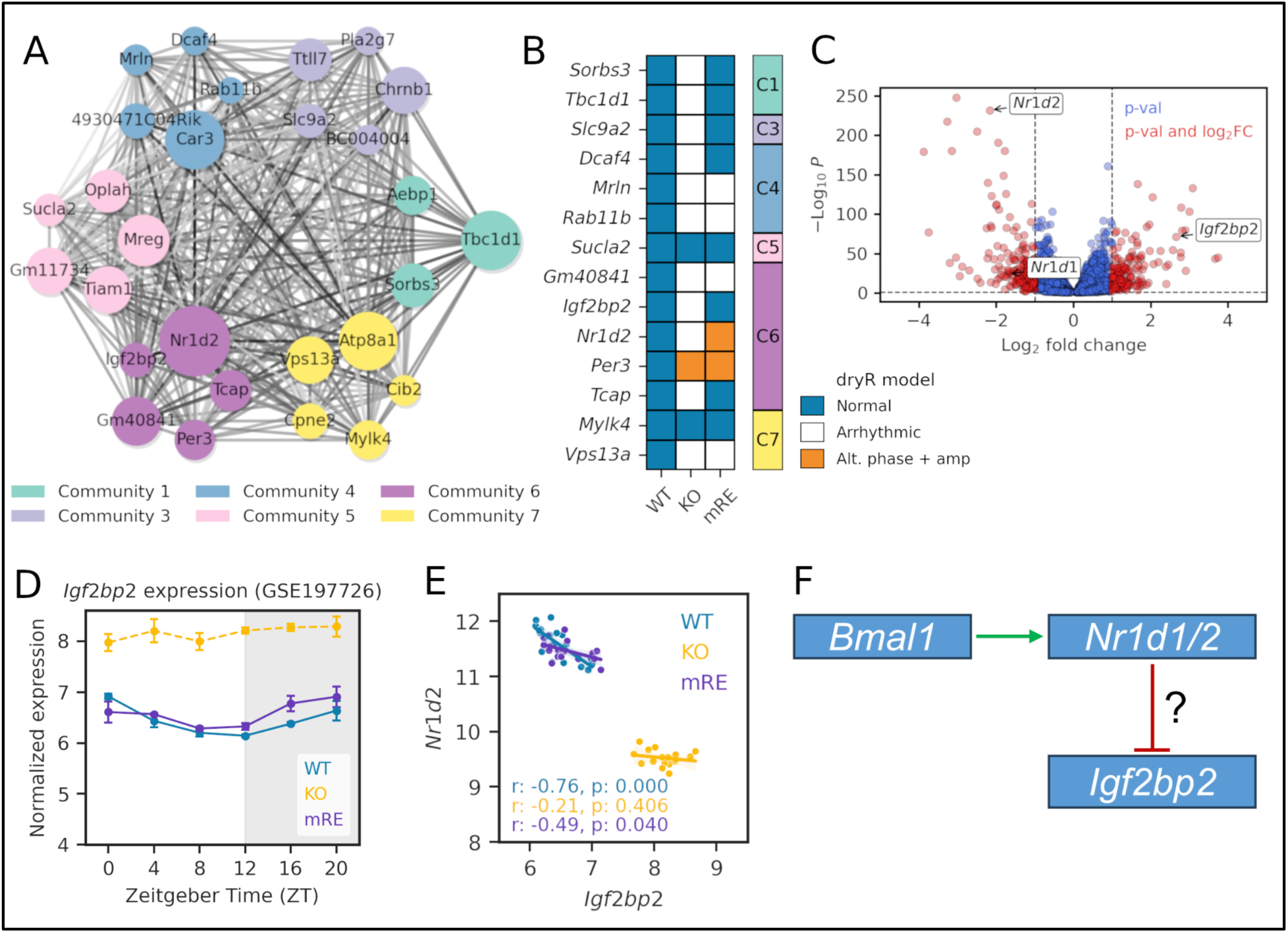
Igf2bp2 is integrated within the circadian gene network and exhibits genotype- and context-dependent rhythmic regulation. **(A)** Communities within the co-expression network of the turquoise module, with node size proportional to weighted degree of hub genes. Darker edge color represents stronger interaction between genes. **(B)** Differential rhythmicity analysis of turquoise module hub genes that were rhythmic in WT genotype in KO and mRE genotypes, highlighting altered oscillatory patterns. **(C)** Volcano plot of differential gene expression identifying significant changes in Nr1d2 and Igf2bp2 in GSE197726 (WT vs KO). **(D)** Gene expression profile of Igf2bp2 across time points in WT, KO, and mRE genotype (GSE197726). Solid lines indicate rhythmic expression, whereas dashed lines indicate arrhythmic expression based on rhythmicity analysis (FDR < 0.10). **(E)** Correlation between Igf2bp2 and Nr1d2 expression in GSE197726 dataset. Pearson correlation coefficients (r) and p-values are shown for each genotype (WT, KO, mRE). **(F)** Proposed Bmal1-Nr1d1/2-Igf2bp2 regulatory axis. Also see Fig. S2, Supplementary Data 1 and Table S1.

To tease apart the relationship between hub genes in the circadian community, we performed differential rhythmicity (dryR) and differential expression analyses. *dryR* analysis (Weger et al., 2021) identified 14 hub genes that were rhythmic in the WT, including the hubs corresponding to the circadian community (*Nr1d2*, *Per3*, *Tcap*, *Igf2bp2*, and *Gm40841*). Notably, four of these five hubs lost rhythmicity in KO but regained it in mRE (Fig. 2A-B). *Nr1d2* was the top hub gene within the circadian community (|GS|: 0.97, |MM|: 0.98, and weighted degree: 46.74), suggesting it plays a key role in the coordinated regulation of genes within this community. The rhythmic oscillation of *Tcap* is regulated by CLOCK:BMAL1 and synergistic interactions with MYOD1 (Hodge et al., 2019; Riley et al., 2022). Hence, the loss of *Tcap* rhythmicity is expected due to loss of *Bmal1* in KO (Fig. 2B), as is disruption of *Per3* rhythmicity (Fig. 2B). However, Gm40841 and insulin-like growth factor 2 binding protein 2 (*Igf2bp2*) have no known circadian function in muscle. IGF2BP2 is an mRNA-binding protein that regulates the translation, transport, and stability of RNAs (Degrauwe et al., 2016). *Igf2bp2* was arrhythmic in KO mice and regained rhythmicity in mRE mice (Fig. 2B). Analysis of daily average expression revealed that *Igf2bp2* was upregulated in KO, suggesting that *Igf2bp2* is an indirect target of BMAL1, since direct targets of BMAL1 - a transcriptional activator - are predicted to be downregulated in KO. *Nr1d2* is a direct target of BMAL1 and, as expected, was downregulated in KO (Fig. 2C). Thus, upregulation of *Igf2bp2* expression in *Bmal1* KO may be due to an indirect mechanism, possibly involving derepression stemming from loss of *Nr1d2* expression in the absence of *Bmal1*. In line with this notion, we observed a strong anti-correlation between *Igf2bp2* and *Nr1d2* expression in WT and mRE mice, whereas the correlation was lost in KO, in addition to loss of *Igf2bp2* rhythmicity (Fig. 2B, 2D, 2E).

In order to understand the key biological processes tied to the *Nr1d2* and *Igf2bp2*, we performed gene ontology on their common neighbors present in the circadian community (see Supplementary Data 1). As expected, terms associated with circadian regulation of gene expression were top enrichments. In addition, processes such as response to insulin (*Gpam, Sesn3, Slc25a33, Cry1* and *Bcar3)* and regulation of protein ubiquitination (*Amer1*, *Cdc14b*, *Nlrc3, Cry1,* and *Per2*) were significant enrichments (see Fig. S2). Collectively, these results from co-expression network analysis suggest that *Igf2bp2* is an important circadian gene in skeletal muscle, which may be a target of *Nr1d2* (Fig. 2F).

### *Igf2bp2* is a clock-controlled gene in skeletal muscle

To validate *Igf2bp2* as a clock-controlled gene, we measured the ZT8 and ZT20 expression of *Bmal1, Nr1d2 (a.k.a., Rev-erbβ),* and *Igf2bp2* in gastrocnemius muscle of WT mice (*Bmal1^wt/wt^*; *Hsa-*Cre^Tg/0^) and muscle-specific *Bmal1* knockout mice (*Bmal1^flox/flox^*; *Hsa-*Cre^Tg/0^), which do not exhibit the shortened lifespan and developmental abnormalities of KO and mRE mice. As expected, *Rev-erbβ* was downregulated in the absence of *Bmal1*. Conversely, *Igf2bp2* was upregulated at ZT8 (Fig. 3A), which coincides with peak *Rev-erb* expression and activity (Gutierrez-Monreal et al., 2020). These results are consistent with co-expression network analyses. To more directly implicate REV-ERB in *Igf2bp2* regulation, we analyzed a publicly available skeletal muscle *Nr1d1/2* double knockout dataset (J. Liu et al., 2025). The relative expression of *Igf2bp2* was significantly higher in *Nr1d1/2* double knockout compared to WT at both ZT10 (log_2_FC of 3.09) and ZT22 (log_2_FC of 3.34) (J. Liu et al., 2025), showing that loss of *Rev-erb* is sufficient to upregulate *Igf2bp2* (Fig. 3B).

**Figure 3.**
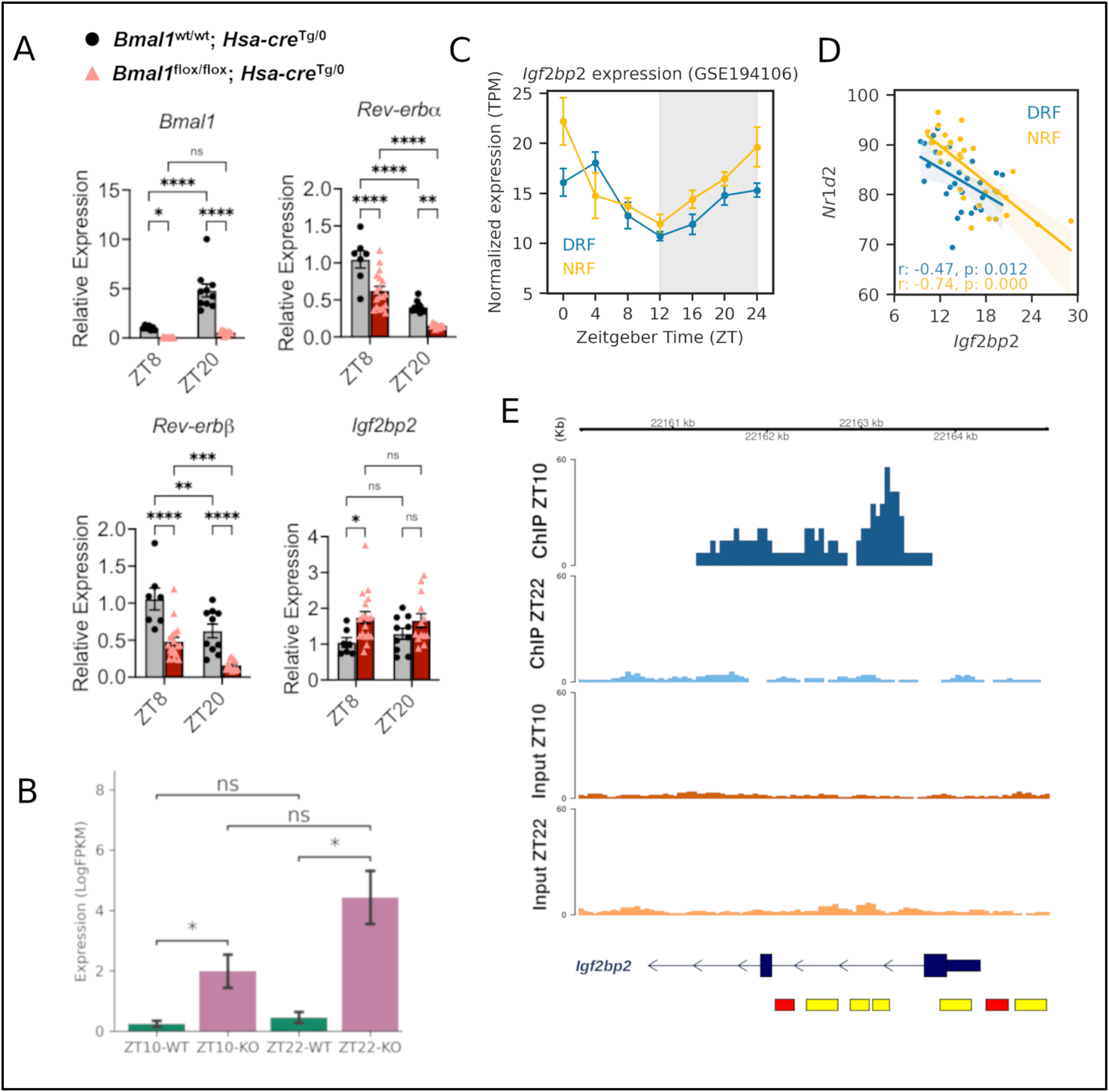
Potential Regulation of Igf2bp2 as a clock-controlled gene. **(A)** Differential regulation of Igf2bp2 and Rev-erb in muscle-specific Bmal1 knockout mice. Gene expression measured by qPCR in gastrocnemius muscle from 16-20-week-old male (M) and female (F) mice housed in standard 12 h light, 12 h dark conditions. ZT - zeitgeber time; ZT0=lights on, ZT12=lights off. Bmal1^wt/wt^; Hsa-cre^Tg/0^ - wild-type (WT); Bmal1^flox/flox^; Hsa-cre^Tg/0^ - muscle-specific Bmal1 knockout (KO). Two-way ANOVA, Fisher’s LSD post hoc test, *p<0.05, **p<0.01, ***p<0.001, ****p<0.0001, ns - not significant, WT ZT8 n=7 (F=4, M=3), WT ZT20 n=10 (F=5, M=5), KO ZT8 n=18 (F=11, M=7), KO ZT20 n=13 (F=7, M=6). **(B)** Expression of Igf2bp2 at ZT10 and ZT22 in Nr1d1/2 DKO relative to WT (P < 0.05) (GSE249730). **(C)** Gene expression profile of Igf2bp2 across time points in different feeding regime (DRF: daytime restricted feeding and NRF: nighttime restricted feeding, GSE194106). Solid lines indicate rhythmic expression based on rhythmicity analysis (FDR < 0.10). Data represented as mean ± SEM. Time is indicated in hours post-synchronization or Zeitgeber time (ZT), as reported in the original study (see Table S2). **(D)** Correlation between Igf2bp2 and Nr1d2 expression in GSE194106. Pearson correlation coefficients (r) and p-values are shown for both feeding regimes. **(E)** ChIP-seq peak of REV-ERBα near the exon 1 of Igf2bp2 gene at ZT10 analyzed from the publicly available dataset GSE263635. The red and yellow rectangles represent the putative promoter and enhancer regions, respectively. Also see Fig. S3-S4, Supplementary Data 2 and Table S2.

Next, to determine whether *Igf2bp2* is commonly identified as a rhythmic gene in muscle, we employed MetaCycle and RAIN to nine published skeletal muscle circadian time-series datasets from mouse and human (see Table S2). We found that *Igf2bp2* was rhythmic and anti-correlated with *Nr1d2* in 90% of WT mice experimental groups encompassing different feeding regimes and ages (Fig. 3C-D, also see Fig. S3-S4). In addition, *Igf2bp2* exhibited rhythmic expression in most of the tissue-specific *Bmal1* reconstituted experimental groups (see Fig. S3-S4). Conversely, all global *Bmal1* KO experimental groups exhibited arrhythmic *Igf2bp2* (see Fig. S3-S4). RAIN predicted a modest oscillation of *Igf2bp2* in several of the human datasets that were analyzed (p < 0.05).

Given the established role of REV-ERBs as transcriptional repressors, we next sought to determine whether REV-ERBβ could bind to the *Igf2bp2* locus in skeletal muscle. Although REV-ERBβ skeletal muscle ChIP-seq datasets are not available, we analyzed the publicly available REV-ERBα skeletal muscle ChIP-seq data, since both REV-ERBα and β bind to the same canonical motif (Guillaumond et al., 2005). REV-ERBα ChIP-seq analysis revealed a significant peak near the exon 1 region of *Igf2bp2* gene at ZT10 but not at ZT22 (Fig. 3E, also see Supplementary Data 2). This peak overlaps with a promoter region of *Igf2bp2* (Fig. 3E). These data suggest that REV-ERB may directly regulate *Igf2bp2* transcription.

### REV-ERBβ represses *Igf2bp2* transcription via GCC motifs in the *Igf2bp2* promoter

Thus far, our findings point to a temporal mechanism in which REV-ERBβ regulates *Igf2bp2* expression. To functionally test whether REV-ERBβ represses *Igf2bp2* expression, the *Igf2bp2* promoter region (-2kb to +500bp relative to TSS) containing the putative REV-ERB binding site was cloned into a luciferase reporter construct (Igf2bp2-WT-luc) (Fig. 4A). Transient transfection experiments in differentiated C2C12 myotubes revealed that REV-ERBβ, but not REV-ERBα, repressed Igf2bp2-WT-luc signal in a dose-dependent manner, by upwards of -67% (Fig. 4B). Notably, sequence analysis of the promoter region did not reveal a canonical ROR-response element within the REV-ERB binding site. However, multiple GCC sequences, which are common in top REV-ERB binding motifs (Fig. 4C), were identified. To determine their involvement, all GCCs within the REV-ERB binding site (-131bp to +183bp relative to TSS) were mutated to ATA (Igf2bp2-Mut-luc) (Fig. 4D). REV-ERBβ was unable to repress Igf2bp2-Mut-luc signal, indicating that the putative REV-ERB regulatory site is functional and that GCC motifs are important for REV-ERBβ-mediated repression of Igf2bp2 transcription (Fig. 4B). Altogether, these results demonstrate repression of *Igf2bp2* transcription by REV-ERBβ and implicate *Igf2bp2* in circadian regulation of muscle metabolism.

**Figure 4.**
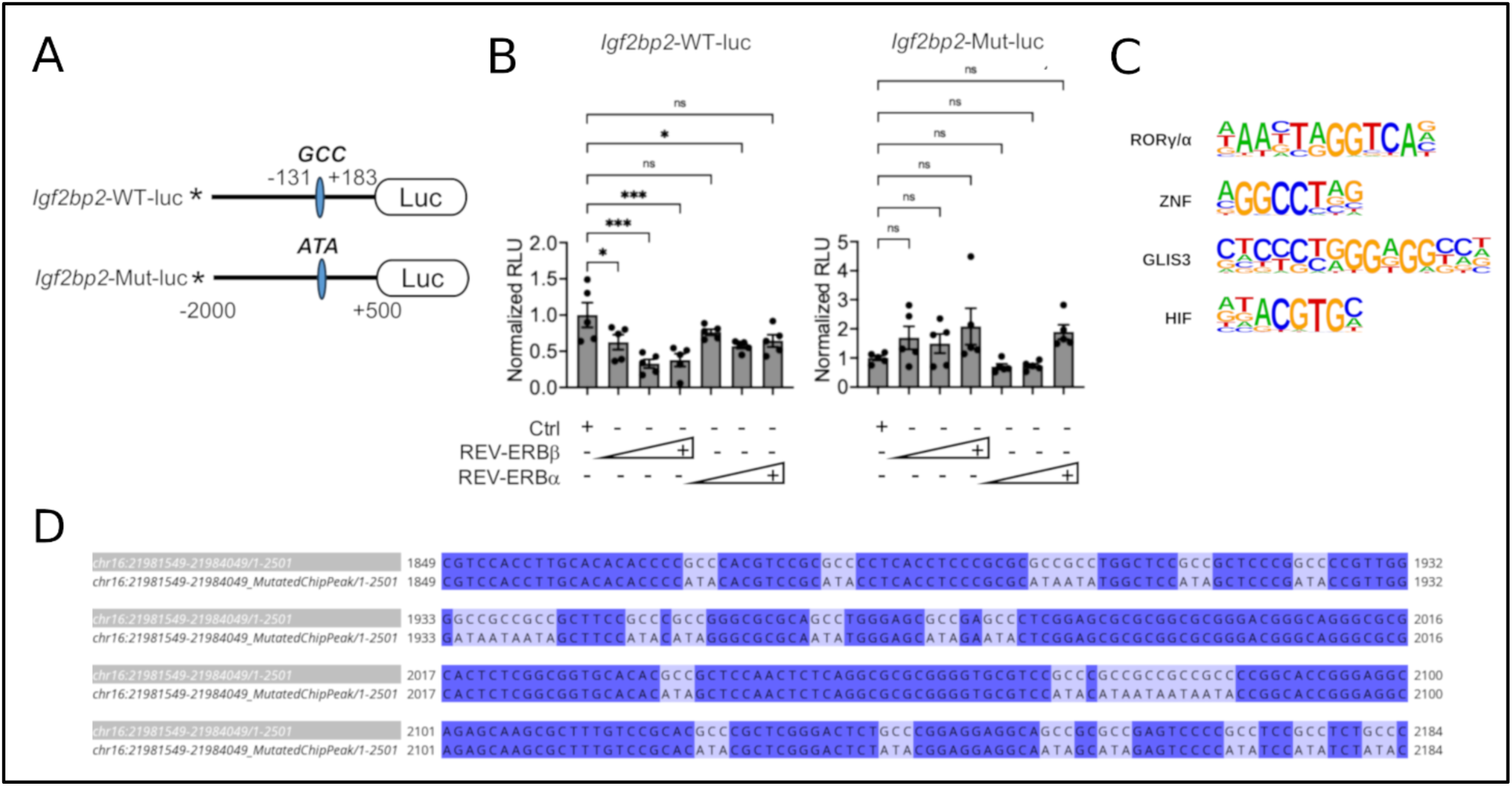
REV-ERBβ Represses Igf2bp2 Transcription Via a GCC Motifs in the Igf2bp2 Promoter. **(A)** Schematic of luciferase reporter constructs containing the Igf2bp2 promoter region. The wild-type construct (WT GCC) harbors the intact REV-ERB response element, while the mutant construct (Mut ATA) contains a disrupted binding motif. Luciferase (luc) reporter driven by the wild-type (WT) Igf2bp2 promoter (-2kb to +500bp relative to TSS) or GCC->ATA mutated (Mut) promoter (-131bp to +183bp relative to TSS). Relative position of Igf2bp2 promoter is indicated. **(B)** Transient transfection assays in C2C12 myotubes. The GCC motif corresponds to a known REV-ERB ChIP-seq peak. REV-ERBβ and REV-ERBα were expressed with CMV-driven vectors at increasing doses of 50, 100, and 200 ng. One-way ANOVA, Dunnett’s post hoc test, *p<0.05, ***p<0.001, ns - not significant, n=5. RLU - relative light unit. **(C)** Enriched REV-ERB binding motifs in the ZT10 ChIP-seq peaks reported in (J. Liu et al., 2025). **(D)** Pairwise alignment between WT and Mut promoter highlighting the GCC->ATA mutations in the REV-ERB binding site (-131bp to +183bp relative to TSS).

## Discussion

Skeletal muscle forms a key component of the peripheral metabolism that adapts to external stimuli such as feeding and exercise. More importantly, the core circadian clock plays a central role in regulating skeletal muscle metabolism and impact of exercise, including glucose utilization, mitochondrial function, and insulin sensitivity. While the disruption of core clock components impair these processes (Chaikin et al., 2025; Chatterjee & Ma, 2016; Dyar et al., 2013; J. Liu et al., 2025; Procopio & Esser, 2025; Viggars et al., 2024), the precise molecular mechanisms underlying these effects are incompletely understood.

Here, we used a co-expression network-based approach to define gene modules in a circadian timepoint-based skeletal muscle dataset across three mice genotypes: WT, global *Bmal1* KO, and muscle *Bmal1* reconstituted (mRE) (GSE197726) (Smith et al., 2023). Our initial analysis revealed distinct clustering of gene expression across genotypes, with a higher variability in mRE. Importantly, the analysis of co-expressed modules showed a distinct association pattern with the genotypes, and could capture gene clusters that were restored in mRE. While few modules showed partial restoration in mRE, others remained dysregulated, consistent with previous reports that reconstitution does not fully recover the WT circadian transcriptome (Smith et al., 2023).

Importantly, our findings revealed that *Nr1d2* (REV-ERBβ), a BMAL1-regulated transcriptional repressor, was the most connected hub in the turquoise module. Further organization of the turquoise module into sub-networks identified a circadian community, which provided a potential link between core clock activity and downstream transcriptional networks. For example, the co-occurrence of *Nr1d2* with PAR-bZip transcription factors (*Dbp, Tef*), and *Igf2bp2* within this community suggested a coordinated regulation through a *Bmal1-Nr1d2* axis. More importantly, among the circadian community hubs, *Igf2bp2* was particularly interesting since it regained rhythmicity in mRE with anti-correlated expression to *Nr1d2* in WT and mRE, but not in KO. Further supported with qPCR experiments in WT and muscle-specific *Bmal1* KO, the observation implicated a *Nr1d2-Igf2bp2* regulatory relationship, which was otherwise not known.

The mechanistic underpinnings of this regulation was further characterized using publicly available REV-ERBα ChIP-seq dataset, revealing a time-dependent binding at the *Igf2bp2* locus, and luciferase reporter assays establishing that REV-ERBβ directly represses *Igf2bp2* promoter activity. Together, these findings establish *Igf2bp2* as a clock-controlled gene being regulated through REV-ERBβ.

Functionally, this novel regulatory relationship could highlight incorporation of clock-regulated post-transcriptional control mechanisms (P. Liu et al., 2024; Parnell et al., 2021). More generally, the RNA-binding proteins such as IGF2BP2 have been implicated in metabolic regulation and muscle homeostasis, and its dysregulation has been associated with metabolic disorders (Dai, 2020; Das et al., 2025; Degrauwe et al., 2016; Regué et al., 2019; Shi & Grifone, 2021).

With no direct evidence of *Igf2bp2* as a clock-controlled gene, the integration of *Igf2bp2* within a circadian gene network could have important implications for skeletal muscle biology. For example, loss of *Igf2bp2* in skeletal muscle has been shown to impair muscle mass accrual (Regué et al., 2019), suggesting a critical role in muscle growth and maintenance. In the context of circadian regulation of skeletal muscle physiology, this raises the possibility that *Igf2bp2* could contribute to time-of-day dependent control of muscle homeostasis, along with REV-ERBs (Z. Li et al., 2012; J. Liu et al., 2025).

Mechanistically, IGF2BP2 is a key reader of m6A-modified transcripts, linking it to epitranscriptomic regulation. Emerging evidence suggests that m6A-mediated RNA regulation is involved in both skeletal muscle function and circadian control of gene expression (L. Chen et al., 2023; Dey & Dey, 2025; Fustin et al., 2013; Hastings, 2013; Wang et al., 2025; Zhong et al., 2018). Thus, circadian regulation of *Igf2bp2* may represent a point of convergence between transcriptional oscillations and m6A-dependent post-transcriptional control. Furthermore, feeding and fasting cycles, which act as dominant zeitgebers for peripheral tissues, may intersect with this regulatory axis (Kumar et al., 2024). Lastly, this axis could also highlight potential therapeutic opportunities through pharmacological modulation of REV-ERB and IGF2BP2 to target metabolic dysfunction arising as a result of disrupted rhythms (Berwanger et al., 2026; Solt et al., 2012; Srivastava et al., 2025). Overall, these observations position REV-ERBβ regulated *Igf2bp2* as a potential mediator linking circadian timing, epitranscriptomic regulation, and skeletal muscle metabolism. However, future circadian studies and multi-omics approaches will be essential to establish their functional consequences in metabolic disease and muscle physiology.

### Limitations of the study

The relationship between metabolism and circadian rhythms is well established in various tissues such as liver, heart and skeletal muscles. However, the precise regulatory mechanisms that govern this crosstalk are still being deciphered. In this work, by inferring gene co-expression networks using bulk RNA sequencing data from specific mice genotypes, we have been able to establish a novel regulatory axis involving *Bmal1, Nr1d2 and Igf2bp2*. We present both computational and experimental evidence for the same. However, there are certain limitations such as- we opted for an all-sample network construction, regardless of time point, to increase statistical power and to assess whether, within the same network framework, changes in co-expressed gene interactions could be observed across genotypes, and identified novel circadian genes. Constructing independent networks based on genotype and circadian time points is an approach that may provide additional insights, but this was not feasible given limited data. In addition, we utilized published REV-ERBα ChIP-seq data to identify potential regulatory sites within the *Igf2bp2* promoter region because REV-ERBβ ChIP-seq data is not currently available. Thus, REV-ERBβ may bind additional *Igf2bp2* promoter or enhancer sites that serve important regulatory functions and have not yet been identified. The results may get stronger evidence in presence of REV-ERBβ ChIP-seq data.

## Resource Availability

### Lead contact

Further information and requests for codes, resources, reagents, should be directed to, and will be fulfilled by, the lead contacts, Dr. Ashutosh Srivastava (codes) and Dr. Kevin B. Koronowski (resources and reagents).

### Materials availability

Requests for materials should be directed to the lead contact, Dr. Kevin B. Koronowski, and will be subject to a materials transfer agreement.

### Data and code availability

All datasets generated and/or analyzed in this study are included in this published article and its supplementary information files, with source data provided. Publicly available datasets used in this study are listed in the Key Resources Table. Any additional information required to reanalyze the data reported in this paper is available from the Lead Contacts.

## Acknowledgements

This project was supported by Department of Biotechnology (DBT), Ministry of Science and Technology, Government of India (A.S.), Council for Scientific and Industrial Research (CSIR), New Delhi, India - 09/1031(13466)/2022-EMR-I (S.S.), Indian Institute of Technology Gandhinagar, Gandhinagar, Gujarat, India (V.A.M.). The Koronowski lab is supported by the National Institute of General Medical Sciences under award number R35GM150618, the Max and Minnie Tomerlin Voelcker Fund, and the American Heart Association (25CDA1451928). This work was supported by the San Antonio Nathan Shock Center of Excellence in the Biology of Aging under award number P30AG013319.

## Author Contributions

A.S., S.S., V.A.M. designed the study and wrote the original manuscript draft. K.B.K and Q.Z. designed, performed, interpreted the experimental data, and wrote the experimental results.

A.S. and K.B.K refined and assisted in manuscript writing. V.A.M. and S.S. performed the computational analysis. A.S. and K.B.K. are the lead contacts. All authors contributed to manuscript editing and approving the final manuscript version.

## Declaration of Interests

The authors declare no competing interests.

## Declaration of Generative AI and AI-assisted technologies in the writing process

During the preparation of this work the authors V.A.M. and S.S. used ChatGPT in order to refine the text. After using this tool/service, all the authors reviewed and edited the content as needed and take full responsibility for the content of the published article.

## STAR★ methods

### Key resources table

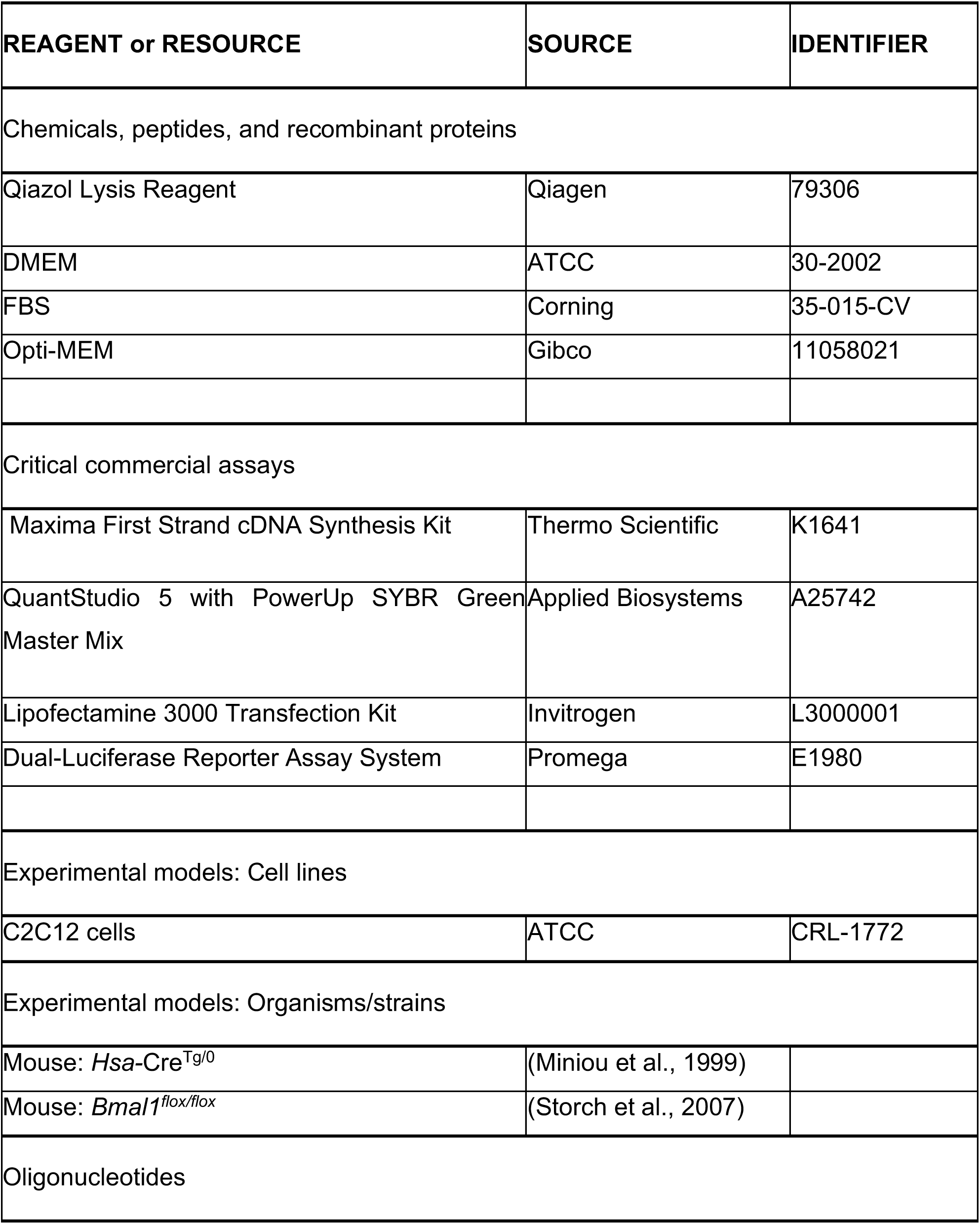
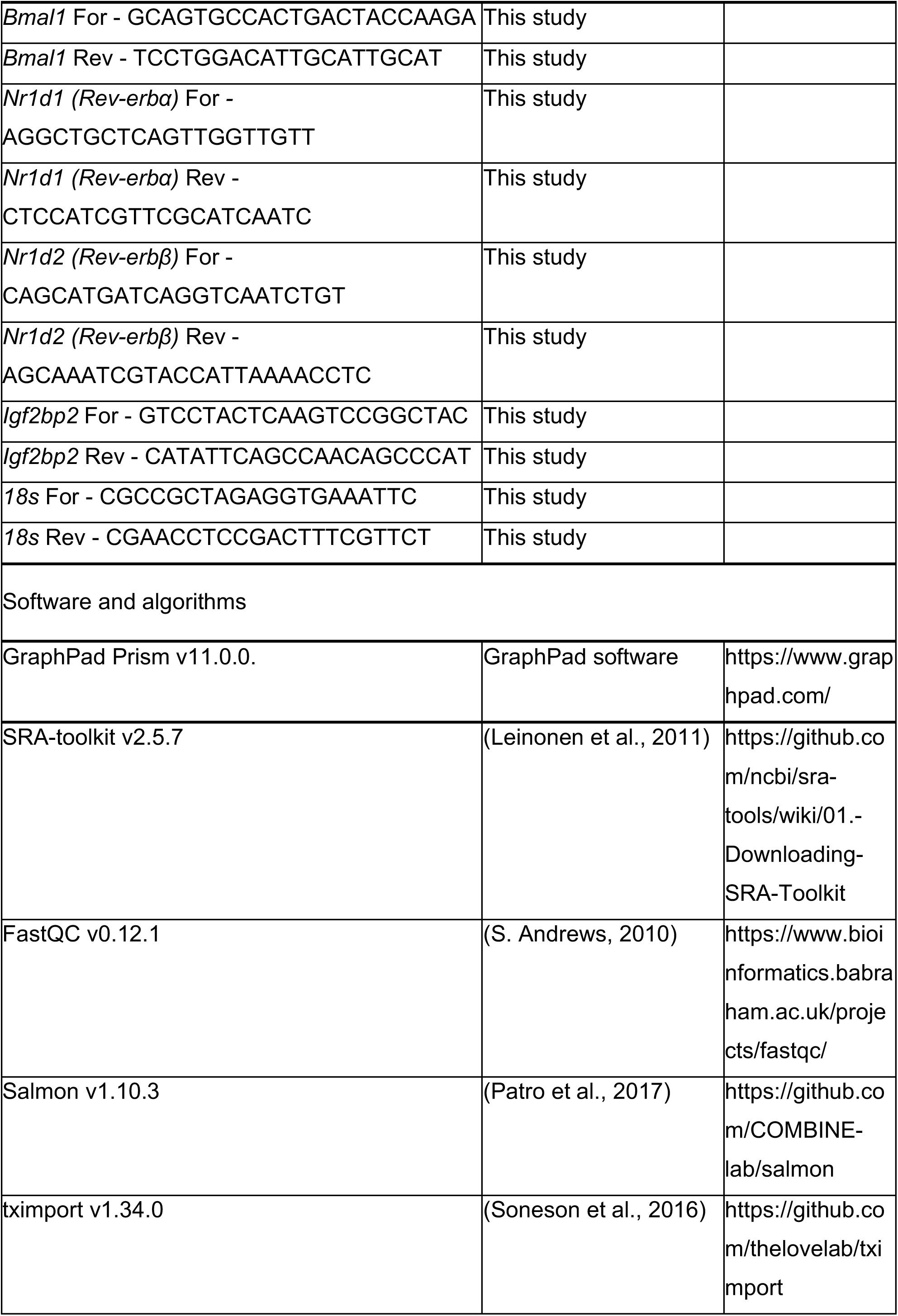
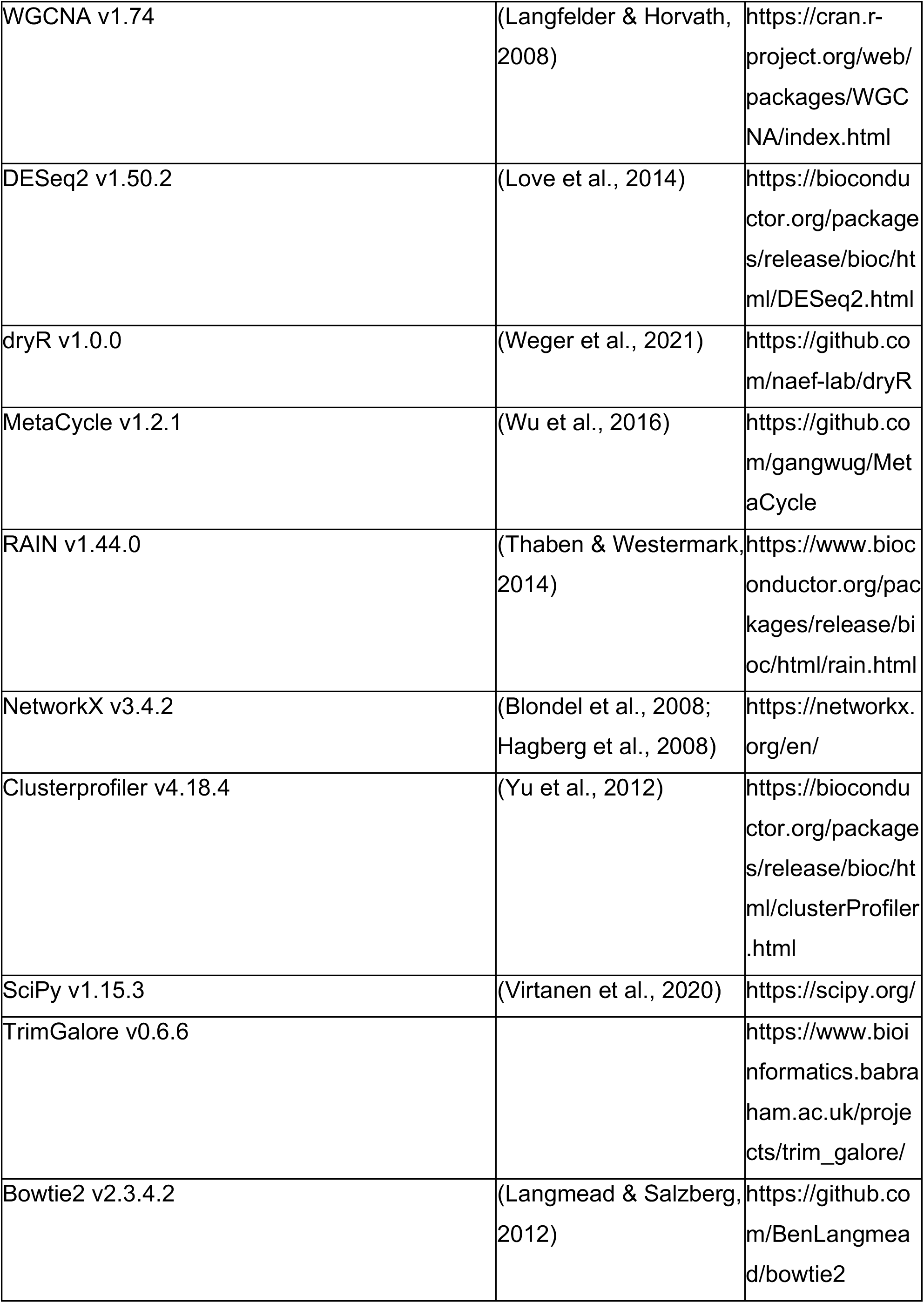
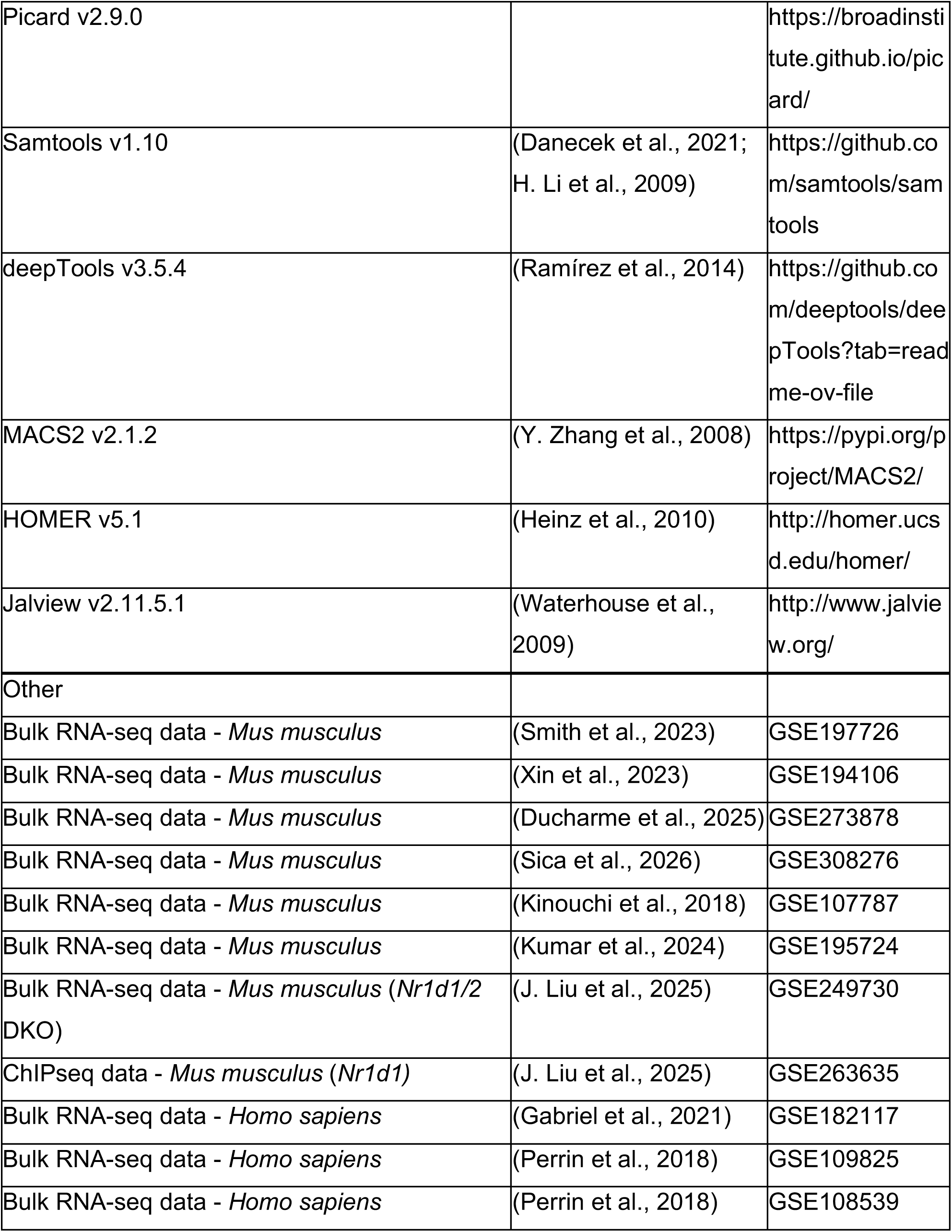

### Experimental Model and Study Participant Details

#### Animals

Animal experiments were conducted at UT Health San Antonio in accordance with the National Research Council’s Guide for the Care and Use of Laboratory Animals. All experiments were conducted with approval from the local Institutional Animal Care and Use Committee and ARRIVE guidelines were followed wherever possible. *Bmal1*-flox mice (Storch et al., 2007) on the C57BL/6J background were crossed with the Hsa-Cre mice (Miniou et al., 1999) to generate mice with muscle-specific knockout of *Bmal1*. Experimental genotypes: 1. Wild-type (WT) - *Bmal1^wt/wt^*; *Hsa-*Cre^Tg/0^. 2 Muscle-specific *Bmal1* knockout (KO) - *Bmal1^flox/flox^*; *Hsa-*Cre^Tg/0^. Male and female mice ages 8-14 weeks were randomly assigned to groups and entrained to standard 12 hr light (∼100 lux), 12 hr dark cycles. Mice were fed *ad libitum* with chow diet (Research Diets, D12450J) for 8 weeks, at which point gastrocnemius muscle was harvested at ZT8 or ZT20 and flash frozen in liquid nitrogen for storage at -80°C.

### Method Details

#### Quantitative PCR analysis

Gastrocnemius muscle was crushed in liquid nitrogen into powder-like consistency using a mortar and pestle. Approximately half of one muscle was homogenized in Qiazol Lysis Reagent (Qiagen, 79306) by vortexing followed by 15 min incubation at room temperature (RT). Chloroform was added to samples at a 1 to 5 ratio and incubated for 15 min at RT. Samples were then centrifuged for 15 min at 12,000 x g and the supernatant was transferred to a new tube. 100% isopropanol was added at a 1 to 1 ratio and samples were incubated for 10 min at RT. Samples were then centrifuged for 15 min at 12,000 x g and the resulting pellet was washed twice with 70% ethanol. The RNA pellet was then resuspended in RNase- and DNase-free water. cDNA was prepared from 1 ug RNA using the Maxima First Strand cDNA Synthesis Kit (Thermo Scientific, K1641). Quantitative real-time PCR was performed on a QuantStudio 5 with PowerUp SYBR Green Master Mix (Applied Biosystems, A25742) and normalized to *18s* control. Primer sequences:

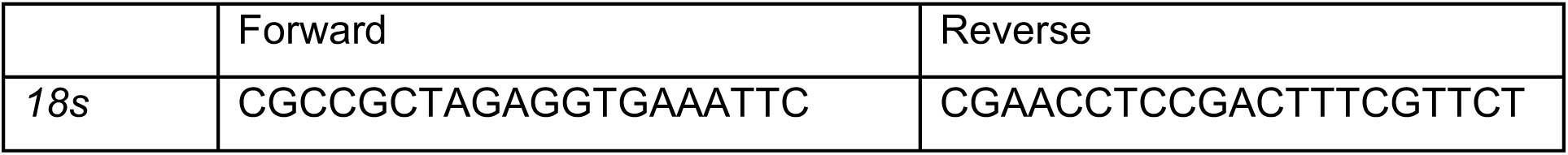
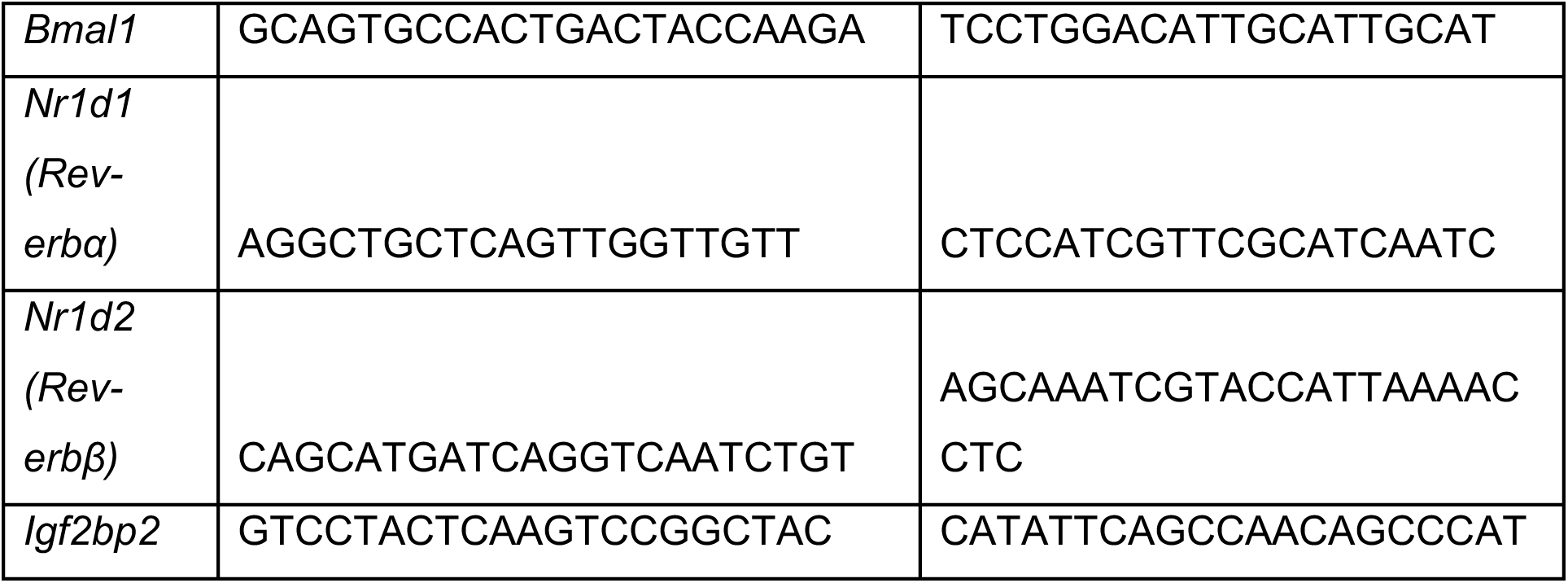

#### Cell culture and luciferase assays

##### C2C12 cells

C2C12 cells (ATCC, CRL-1772) were cultured in maintenance medium consisting of DMEM (ATCC, 30-2002) supplemented with 10% FBS (Corning, 35-015-CV) and 1% penicillin-streptomycin and were kept from expanding past 80% confluency. For experiments, cells were plated at confluency in 96-well plate and reverse transfected at the time of plating with varying doses of the respective plasmids according to the Lipofectamine 3000 Transfection Kit (Invitrogen, L3000001) by 4-h incubation with packaged particles in Opti-MEM (Gibco, 11058021). Following the 4-h transfection, cells were washed 3 x with PBS and differentiated into myotubes for 3 days using differentiation medium consisting of DMEM supplemented with 2% Horse serum and 1% penicillin-streptomycin, which was changed daily, leaving behind 20% of the media with each media change. After differentiation, luciferase activity was measured using the Dual-Luciferase Reporter Assay System (Promega, E1980), according to the manufacturer’s protocol, in a Varioskan LUX Multimode Microplate Reader (Thermo Fisher Scientific). Firefly luciferase signal from the experimental luciferase reporter plasmid was normalized to the control *Renilla* luciferase plasmid signal.

#### Design of Luciferase Reporter Constructs

Putative promoter regions of *Mus musculus Igf2bp2* were identified using Ensembl (genome build GRCm39). Coordinates were extracted relative to the GRCm39 assembly. As *Igf2bp2* is encoded on the reverse strand, all promoter sequences were reverse complemented prior to cloning, and reported here in genomic coordinate space (GRCm39). *Samtools* command line package was used to extract the promoter sequence (Danecek et al., 2021; H. Li et al., 2009). The extracted sequence was then reverse complemented with the *Seqtk* package.

We took -2kb to +500 region relative to the TSS of *Igf2bp2*. The following are the ChIP-seq peak region for *Nr1d1* near *Igf2bp2* and the overlapping region of this peak and the putative promoter region. *Nr1d1* ChIP-seq peak coordinates: chr16:21,981,861-21,982,187 (GRCm39). Overlapping region of the ChIP-seq peak and the putative promoter region: chr16:21,982,039-21,982,187 (GRCm39).

The *Igf2bp2* promoter sequence (reverse complemented) was used to mutate the overlapping regions in the promoter sequence. The GCC motifs were enriched in the putative promoter region of *Igf2bp2* and have been reported in REV-ERB binding motifs, such as ZNF and GLIS3 (J. Liu et al., 2025). Based on this rationale, the GCC motifs were selected for targeted mutations and were replaced with ATA. Pairwise-alignment for WT and Mut sequences was performed with Jalview (v2.11.5.1) (Waterhouse et al., 2009).

Plasmids were generated by VectorBuilder (Chicago, IL, USA). The mouse Nr1d1 (NM_145434.4) or Nr1d2 (NM_001425894.1) coding sequence was cloned into a mammalian expression vector under the control of the CMV promoter: pRP[Exp]-EGFP-CMV>mNr1d1[NM_145434.4] (REV-ERBα); pRP[Exp]-EGFP-CMV>mNr1d2 [NM_001425894.1] (REV-ERBβ). Expressed proteins do not include C- or N-terminal tags. For reporter plasmids, wild-type (WT) or mutated (Mut) *Igf2bp2* promoter sequences were synthesized and inserted to drive expression of firefly luciferase: pRP[Pro]-hRluc-{Igf2bp2 WT}>Luciferase (*Igf2bp2*-WT-luc); pRP[Pro]-hRluc-{Igf2bp2 Mutated}>Luciferase (*Igf2bp2*-Mut-luc). Plasmids were validated by restriction enzyme digestion. Plasmid maps and sequences are available as supplemental files. Plasmids are available upon request.

#### Curation of skeletal muscle circadian time-series data

The publicly available circadian RNA-seq datasets were obtained from Gene Expression Omnibus (GEO) (Barrett et al., 2005). Briefly, the NCBI GEO query *“(circadian[All Fields] OR ZT[All Fields]) AND "skeletal muscle"[All Fields] AND ("Mus musculus"[porgn] OR "Homo sapiens"[porgn]) AND "Expression profiling by high throughput sequencing"[Filter]”* was used to search for circadian time-series datasets. The *Series* and *Tissues* filters were applied to refine the results and circadian time-series skeletal muscle datasets were considered. Based on the criteria, the GEO dataset GSE197726 was selected for the analysis (Smith et al., 2023). The bulk RNA-seq dataset comprised gastrocnemius muscle biopsies collected from wild-type (hereafter referred to as WT), whole body *Bmal1* knockout (hereafter referred to as KO), and skeletal muscle *Bmal1* reconstituted (hereafter referred to as mRE) mice every 4 h from Zeitgeber Time ZT0-ZT20 (n = 3 per time point, total 54 samples with 18 per genotype). The selection of this dataset was based on the three different genotypes that could help investigate the regulation by circadian clock in presence of standard experimental conditions. In addition, other publicly available circadian time-series datasets from human and mouse skeletal muscles biopsies/primary myotubes were analyzed for expression of *Igf2bp2*, *Nr1d2*, and *Bmal1*. The details of all the datasets analyzed are summarized in Table S2.

#### RNA-seq data processing and quality control

All the raw sequencing files (SRA format) corresponding to GSE197726 were downloaded from the NCBI Sequence Read Archive database using the *prefetch* function within the SRA-toolkit (Version 2.5.7) (Leinonen et al., 2011; The SRA Toolkit Development Team, 2025). All the SRA files were converted to Fastq format using the *fasterq-dump* function. Quality control of all the Fastq files was carried out using FastQC package (Version 0.12.1) (S. Andrews, 2010). All libraries showed per-base sequence quality and little to no adapter content and hence no trimming procedures were performed. The duplicated and over-represented sequences were not removed.

Salmon (Version 1.10.3) was used for transcript quantification in the mapping-based mode (Patro et al., 2017). A decoy-aware transcriptome index was built using the mouse transcriptome version M36 and the genome sequence primary assembly version GRCm39 from GENCODE by following the recommended workflow in Salmon documentation (Mudge et al., 2025). Salmon quant command was used to quantify transcripts against the index.

Transcript-level quantifications were summarized to gene-level counts using the tximport R package v1.34.0 (Soneson et al., 2016). Ensembl gene annotation for Mus musculus (Ensembl release 113; EnsDb database, AnnotationHub record AH119358) was retrieved with Annotationhub, and transcript-to-gene mapping (tx_id to gene_id) were constructed. Salmon quantifications were imported from the quantification files provided by salmon using the *tximport* function and summarized at the gene level. The gene-level counts were then extracted from the output of *tximport* function.

#### Data preprocessing and exploratory data analysis

Gene and sample-level quality control was performed using the *goodSampleGenes* function implemented in the WGCNA package (Langfelder & Horvath, 2008). The genes that were flagged as unreliable were filtered out from further analysis. The raw gene expression matrix was then converted into a DESeq2 dataset using *DESeqDataSetFromMatrix* without specifying any design formula (Love et al., 2014). Next, genes with less than 15 counts in more than 75% of samples (41/54) were excluded from the analysis. Lastly, variance stabilization was performed using the *vst* function (variance stabilizing transformation), implemented in DESeq2 (Love et al., 2014). The final dataset consisted of a normalized gene expression matrix for 13918 genes and 54 samples (see Fig. S1).

To understand the gene expression patterns associated with each genotype, hierarchical clustering and principal component analysis (PCA) were performed on the variance-stabilized counts matrix. Briefly, hierarchical clustering was performed using the average linkage method based on the Euclidean distances. The distance matrix was used to generate a dendrogram with the *hclust* function in R. PCA was performed using the *prcomp* function in R (R Core Team, 2021).

#### Differential gene expression analysis

Raw gene-level count matrices for WT and KO samples (GSE197726) used for WGCNA based network construction were used for differential gene expression analysis. Differential gene expression was performed using the DESeq2 package (Love et al., 2014). A *colData* column was constructed by specifying the sample genotypes (WT vs KO) and time points (Zeitgeber time, ZT). The design of the formula was specified as ∼condition and time-points. Genes with adjusted p-value < 0.05 and |log₂ fold change| ≥ 1 were considered significantly differentially expressed.

Another publicly available skeletal muscle *Nr1d1/2* double knockout dataset, GSE249730 (hereafter referred to as DKO), was used to investigate the expression levels of *Igf2bp2* (J. Liu et al., 2025). The dataset consisted of skeletal muscle RNA-seq data for Wiltype (WT) and DKO at ZT10 and ZT22 (3 replicates per timepoint per genotype, resulting in 12 samples). The FPKM values from this data were used to visualize the expression of *Igf2bp2*. The results table from the differential gene expression analysis reported in the original paper was examined to extract the fold change values of *Igf2bp2* (J. Liu et al., 2025).

#### Differential rhythmicity and circadian rhythmicity detection

The DryR R package was used to identify oscillating genes. The analysis was performed with the *dryseq* function within the DryR package, specifying experimental group labels (WT, KO, mRE) and corresponding time points (ZT0, ZT4, ZT8, ZT12, ZT16, ZT20; three replicates per condition. The filtered gene set used for WGCNA was used for DryR analysis. However, it should be noted that after converting the Ensembl IDs of the 13918 genes to gene symbols only 12619 were parsed into the *dryseq* function (Weger et al., 2021).

Rhythmic gene expression for the genes in GSE197726 was assessed using the MetaCycle package (Wu et al., 2016) and RAIN (Rhythmicity Analysis Incorporating Nonparametric methods) (Thaben & Westermark, 2014).

Rhythmicity using MetaCycle was assessed using the *meta2d* function and integrated methods ARSER, JTK_CYCLE, and Lomb-Scargle. The period search range was set between 20-28 hours. Combined p-values were calculated using Fisher’s method, and Benjamini-Hochberg adjustment was applied to identify significantly oscillating genes (FDR < 0.1) (Wu et al., 2016). The parameters for other experiments such as time and number of samples were adjusted as reported in the original paper.

The RAIN analysis was performed with parameters deltat = 4, period = 24, and nr.series = 3, using the “independent” method to account for repeated measurements. Benjamini-Hochberg correction was applied to RAIN p-values, and genes with FDR < 0.1 were considered rhythmic (Thaben & Westermark, 2014). In addition, the parameters of other datasets for the number of replicates (nr.series) and method (“independent” or “longitudinal”) were set based on the experimental setup and were analyzed independently. The genes crossing the threshold of FDR < 0.1 in either of the algorithms were considered rhythmic.

#### Weighted Gene Correlation Network Analysis (WGCNA)

The WGCNA R package was used to construct the weighted gene co-expression network (Langfelder & Horvath, 2008). The variance-stabilized counts matrix was used to perform weighted gene co-expression network analysis. Firstly, the *pickSoftThreshold* function was used to choose an appropriate power to raise the correlation values, such that the resultant network satisfies the scale-free criterion (Barabási & Albert, 1999). The correlation values are raised to a power in order to emphasize the stronger correlations. A soft power of 18 was chosen to fulfil the scale-free criterion.

The *blockwiseModules* function implemented in the WGCNA package was used for network construction and identification of modules. Each co-expression module represents a subnetwork of highly interconnected genes. The following parameters were used in the *blockwiseModules* function: power = 18, maxBlockSize = 14000, networkType = "unsigned", power = soft_power, minModuleSize = 50, mergeCutHeight = 0.25, numericLabels = FALSE, randomSeed = 1234, verbose = 3.

#### Downstream analysis of WGCNA modules

To obtain the representative expression profile of the genes within each module, the module eigengenes were calculated using the *moduleEigengenes* function implemented in WGCNA. Mathematically, these eigengenes represent the first principal component for the genes per module in all the samples. Next, to explore the association between the co-expression modules and the genotypes, we categorized the genotypes by using one-hot encoding, generating three binary variables: WT, KO, and mRE. Each variable was assigned a value of 1 if the sample belonged to the corresponding genotype and 0 otherwise. Pearson correlation coefficients were then calculated between the module eigengenes and the encoded genotype variables. For the module eigengenes displaying differential correlation patterns with the genotypes, we calculated Gene Significance (GS) and Module Membership (MM) to compute the association of individual genes with the genotype.

#### Network analysis of WGCNA modules

The networks of the selected modules were exported as an edgelist using the *exportNetworkToCytoscape* (TOM threshold = 0.05) provided in the WGCNA package (Langfelder & Horvath, 2008). The degree of each node is the number of connections a node

has in a network. The weighted degree was calculated for each gene. Based on their weighted degree, top 5 percentile genes with |GS & MM| values ≥ 0.5 were defined as hub genes. We identified communities to investigate if the genes involved in similar biological processes were clustered as communities within the module networks. The community structure was determined using the Louvain algorithm as implemented in the *community* subpackage in Networkx (Blondel et al., 2008; Hagberg et al., 2008). The resolution parameter was set to 1.1 to get a good partition of the network. Gene ontology was performed on the turquoise module nodes that interacted with both *Igf2bp2* and *Nr1d2* to identify the functionally associated processes. Gene ontology analysis was performed using *Clusterprofiler* R package (Yu et al., 2012).

#### Gene expression correlation in skeletal muscle circadian datasets

To investigate the relationship between the expression of *Igf2bp2* (IGF2BP2 in humans) and *Nr1d2* ((NR1D2 in humans), pairwise correlations were computed between the two genes across all the datasets and experimental groups. Correlations and corresponding p-values were calculated using *pearsonr* function implemented in Python SciPy library (Virtanen et al., 2020).

#### ChIP-seq data analysis

Publicly available ChIP-seq data for NR1D1 at ZT10 and ZT22 from GSE262635 was analysed using the pipeline described in the original paper (J. Liu et al., 2025). Briefly, raw Fastq files were assessed for quality using *Fastqc*, and low quality bases and adapters were trimmed using TrimGalore (v0.6.6). The filtered reads were aligned to the mm10 mouse reference genome using Bowtie2 (v2.3.4.2) (Langmead & Salzberg, 2012). Duplicates were marked and removed using Picard (v2.9.0), and alignment files were processed with Samtools (v1.10) (Danecek et al., 2021; H. Li et al., 2009). Read coverage normalization and bigWig file generation were carried out using deepTools (v3.5.4) (Ramírez et al., 2014). Peak calling was performed with MACS2 (v2.1.2) (Y. Zhang et al., 2008). All the parameters were kept the same as reported in the original paper (J. Liu et al., 2025). Peak annotations were performed using *annotatePeaks.pl* script implemented in HOMER (v5.1) (Heinz et al., 2010). The ChIP-seq peak region was identified from chr16:22,163,111 - 22,163,437 (mm10 genome reference).

## Quantification and Statistical Analysis

Statistical tests for experiments were performed in Prism (GraphPad Software, Inc.). For comparisons across multiple groups in qPCR experiments, statistical significance was determined by two-way ANOVA and Fisher’s LSD post hoc test. For luciferase reporter studies, statistical significance was determined by One-way ANOVA and Dunnett’s post hoc test. For all figures, * denotes p < 0.05, ** denotes p < 0.01, *** denotes p < 0.001, ****\ denotes p <0.0001, and ns indicates not significant. The statistical significance for each figure is provided in the figure legends.

## Supplemental Information

Document S1: Figures S1-S4, Tables S1-S2, and supplemental references. Related to Figure 1 and 2.

Supplementary Data 1: List of gene modules and communities. Related to Figure 1 and 2.

Supplementary Data 2: MACS peak calls for *Nr1d1* at ZT10/ZT22, and *Nr1d1* ZT10 peak annotation file. Related to Figure 3 and 4.

## Notes

### Competing Interest Statement

The authors have declared no competing interest.

