## Supplementary File for "Co-expression network analysis reveals repression of *Igf2bp2* by REV-ERBβ in skeletal muscle"

### Supplemental PDF

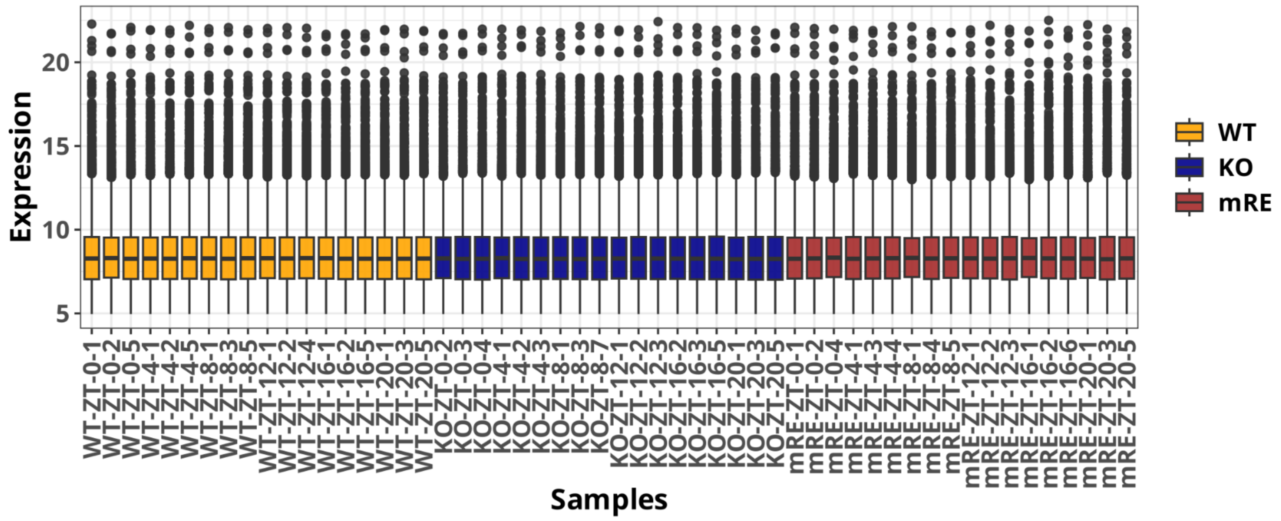


**Figure S1. Variance-stabilizing transformation (VST)-normalized gene expression profile of skeletal muscle circadian time-series RNA-seq data for WT, KO, and mRE genotypes (GSE197726), related to Fig. 1A-B.**

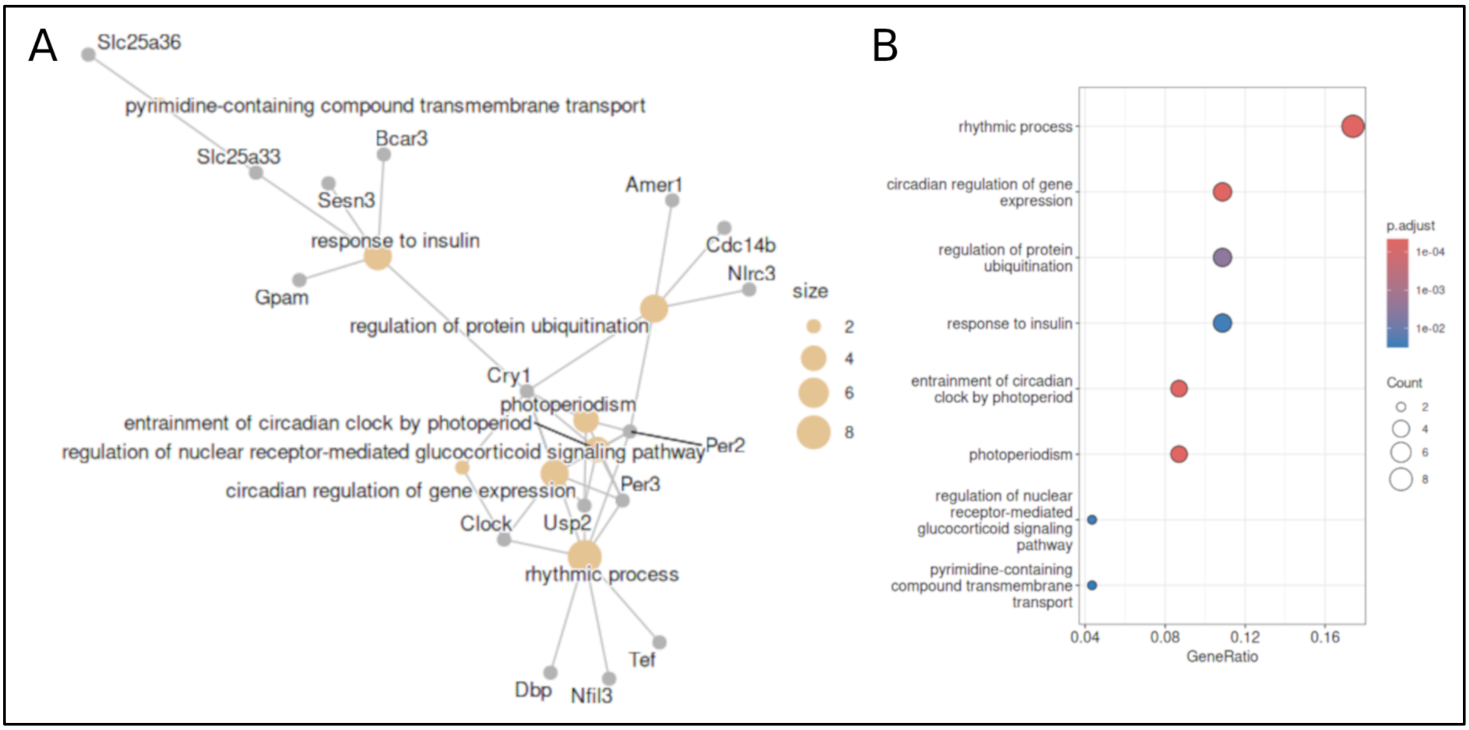


**Figure S2. Gene ontology enrichment analysis of biological processes for common neighbors of *Nr1d2* and *Igf2bp2* present within the circadian community, (A)** Network highlighting the relationship between genes and their associated GO: biological processes. Grey and yellow nodes represent genes and biological processes, respectively. Node size for GO: biological processes highlights the number of genes associated with the term. **(B)** Dot plot highlighting significantly enriched GO: biological processes. Node size and color represent the number of genes associated with the term and the adjusted p-value, respectively. **Related to Fig. 2A-B.**

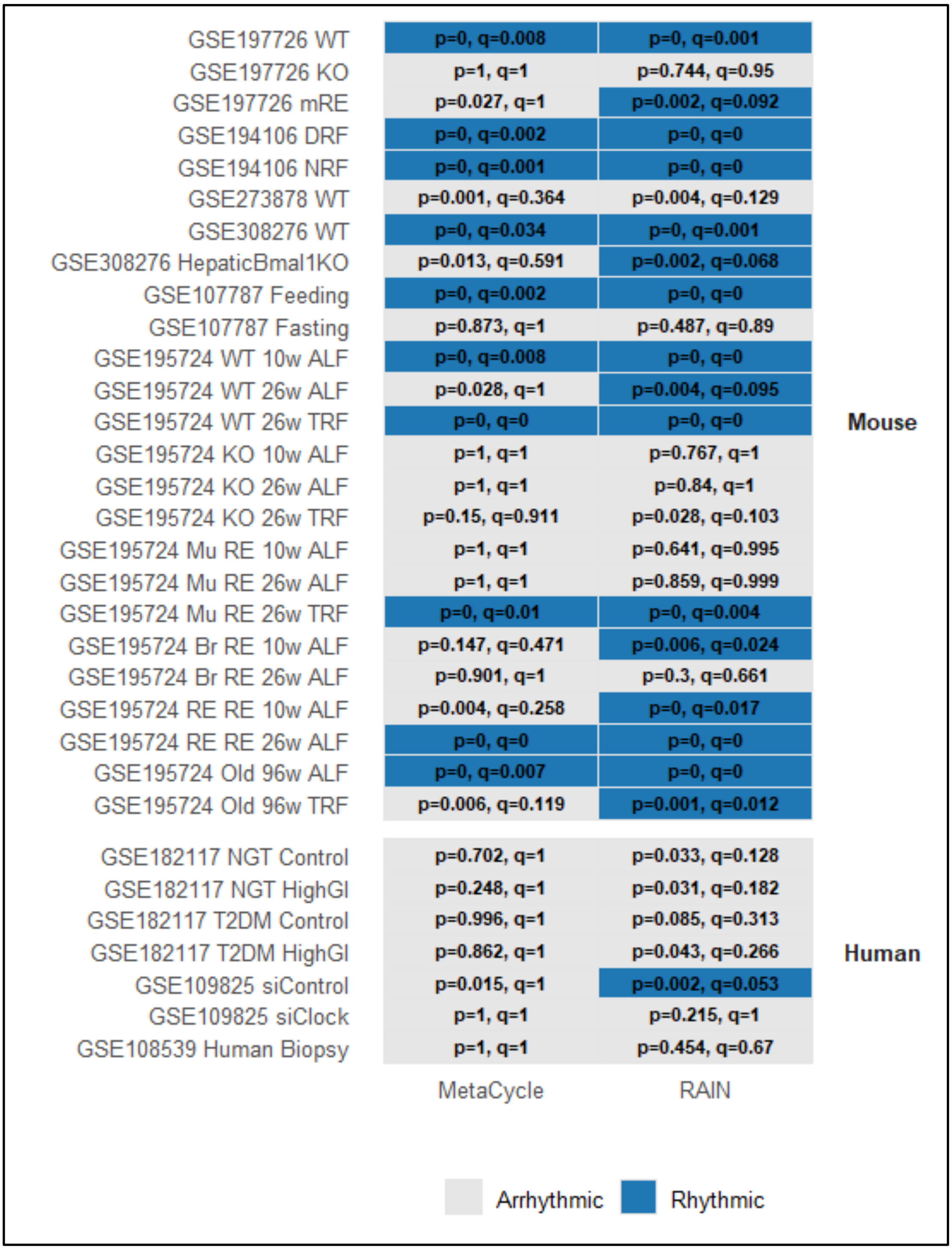


**Figure S3. Rhythmicity of *Igf2bp2* (*IGF2BP2* in human) across publicly available circadian transcriptomic datasets, related to Fig. 2A-B, G-J.**

Rhythmicity of gene expression across multiple mouse and human time-series datasets using two independent algorithms, MetaCycle and RAIN. Each row corresponds to an experimental condition in a dataset, and each column corresponds to the rhythmicity detection method. Rhythmicity was defined as FDR < 0.1 (q).

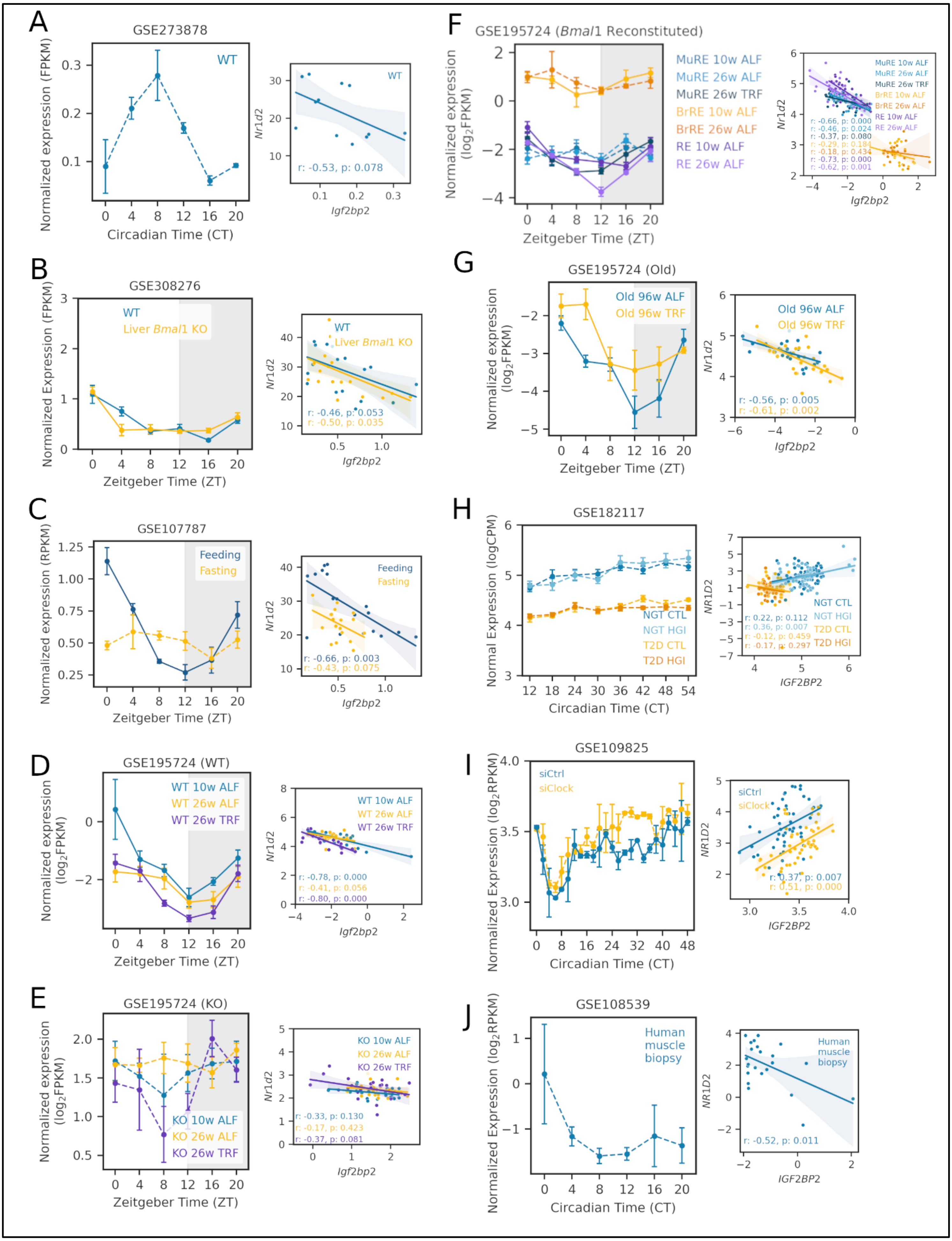
A
GSE273878
WT
Normalized expression (FPKM)
Circadian Time (CT)
Nr1d2
Igf2bp2
r: -0.53, p: 0.078
B
GSE308276
WT
Liver Bmal1 KO
Normalized Expression (FPKM)
Zeitgeber Time (ZT)
r: -0.46, p: 0.053
r: -0.50, p: 0.035
C
GSE107787
Feeding
Fasting
Normalized expression (RPKM)
Zeitgeber Time (ZT)
r: -0.66, p: 0.003
r: -0.43, p: 0.075
D
GSE195724 (WT)
WT 10w ALF
WT 26w ALF
WT 26w TRF
Normalized expression (log2FPKM)
Zeitgeber Time (ZT)
r: -0.78, p: 0.000
r: -0.41, p: 0.056
r: -0.80, p: 0.000
E
GSE195724 (KO)
KO 10w ALF
KO 26w ALF
KO 26w TRF
Normalized expression (log2FPKM)
Zeitgeber Time (ZT)
r: -0.33, p: 0.130
r: -0.17, p: 0.423
r: -0.37, p: 0.081
F
GSE195724 (Bmal1 Reconstituted)
MuRE 10w ALF
MuRE 26w ALF
MuRE 26w TRF
BrRE 10w ALF
BrRE 26w ALF
RE 10w ALF
RE 26w ALF
Normalized expression (log2FPKM)
Zeitgeber Time (ZT)
r: -0.66, p: 0.000
r: -0.46, p: 0.024
r: -0.37, p: 0.080
r: -0.29, p: 0.184
r: -0.18, p: 0.434
r: -0.73, p: 0.000
r: -0.62, p: 0.001
G
GSE195724 (Old)
Old 96w ALF
Old 96w TRF
Normalized expression (log2FPKM)
Zeitgeber Time (ZT)
r: -0.56, p: 0.005
r: -0.61, p: 0.002
H
GSE182117
NGT CTL
NGT HGI
T2D CTL
T2D HGI
Normal Expression (logCPM)
Circadian Time (CT)
NR1D2
IGF2BP2
r: 0.22, p: 0.112
r: 0.36, p: 0.007
r: -0.12, p: 0.459
r: -0.17, p: 0.297
I
GSE109825
siCtrl
siClock
Normalized Expression (log2RPKM)
Circadian Time (CT)
r: 0.37, p: 0.007
r: 0.51, p: 0.000
J
GSE108539
Human muscle biopsy
Normalized Expression (log2RPKM)
Circadian Time (CT)
r: -0.52, p: 0.011

**Figure S4. Gene expression of *Igf2bp2* (*IGF2BP2* in humans) and its association with *Nr1d2* (*NR1D2* in humans) across publicly available circadian transcriptomic datasets, related to Fig. 2D-F and Fig. 3.**

**(A-G)** *Igf2bp2* gene expression across time points in mice skeletal muscle from different datasets and experimental conditions: **(A)** GSE273878 (WT: wild-type), **(B)** GSE308276 (WT: wild-type and Liver *Bmal1* Knockout), **(C)** GSE107787 (Feeding and Fasting regime), and **(D-G)** GSE195724. (different genotypes, ages, and feeding regimes). **(D)** GSE195724-WT (Wild-type, 10 weeks and 26 weeks of age, *ad libitum* and time-restricted feeding). **(E)** GSE195724-KO (Global *Bmal1* knockout, 10 weeks and 26 weeks of age, *ad libitum* and time-restricted feeding). **(F)** GSE195724-RE (only muscle reconstituted *Bmal1* (MuRE), only brain reconstituted *Bmal1* (BrRE), both brain and muscle reconstituted (RE), 10 weeks and 26 weeks of age, *ad libitum* and time-restricted feeding). **(G)** GSE195724-Old (Old mice, 96 weeks of age, *ad libitum* and time-restricted feeding). The right panel shows the correlation between *Igf2bp2* and *Nr1d2,* with Pearson correlation coefficients (r) and p-values indicated.

**(H)** Expression profile of *IGF2BP2* across circadian time in human skeletal muscle samples from individuals with normal glucose tolerance (NGT) or type 2 diabetes (T2D) under control (CTL) or high glucose/insulin (HGI) conditions (GSE182117). **(I)** Gene expression profile of *IGF2BP2* following *CLOCK* knockdown (siClock) compared with control (siCtrl) (GSE109825). **(J)** Gene expression profile of *IGF2BP2* across circadian time points in human skeletal muscle biopsies (GSE108539). The right panel shows the correlation between *IGF2BP2* and *NR1D2* expression, with Pearson correlation coefficients (r) and p-values indicated.

For all time-series expression plots, solid lines indicate rhythmic expression and dashed lines indicate arrhythmic expression based on rhythmicity analysis (FDR < 0.10). Data represented as mean ± SEM. Time is reported as circadian time (CT) or Zeitgeber time (ZT), as reported in original studies.

**Table S1: Hub genes identified within KO genotype based on gene significance, module membership, and weighted network degree in the turquoise module**

| Symbol | GS.genotype_KO | p.GS.genotype_KO | MM.turquoise | p.MM.turquoise | Connectivity | Community |
|---|---|---|---|---|---|---|
| *Tbc1d1* | -0.96 | 0.00E+00 | -0.96 | 0.00E+00 | 38.59 | Community_1 |
| *Sorbs3* | -0.94 | 0.00E+00 | -0.94 | 0.00E+00 | 29.6 | Community_1 |
| *Aebp1* | -0.94 | 0.00E+00 | -0.95 | 0.00E+00 | 27.32 | Community_1 |
| *Chrnb1* | 0.96 | 0.00E+00 | 0.96 | 0.00E+00 | 31.13 | Community_3 |
| *Ttll7* | 0.95 | 0.00E+00 | 0.96 | 0.00E+00 | 29.03 | Community_3 |
| *Slc9a2* | 0.94 | 0.00E+00 | 0.96 | 0.00E+00 | 25.26 | Community_3 |
| BC004004 | 0.94 | 0.00E+00 | 0.96 | 0.00E+00 | 23.93 | Community_3 |
| *Pla2g7* | 0.94 | 0.00E+00 | 0.95 | 0.00E+00 | 23.34 | Community_3 |
| *Car3* | -0.98 | 0.00E+00 | -0.97 | 0.00E+00 | 38.31 | Community_4 |
| 4930471C04Rik | 0.97 | 0.00E+00 | 0.95 | 0.00E+00 | 25.77 | Community_4 |
| *Mrln* | 0.95 | 0.00E+00 | 0.95 | 0.00E+00 | 24.16 | Community_4 |
| *Dcaf4* | 0.95 | 0.00E+00 | 0.95 | 0.00E+00 | 23.4 | Community_4 |
| *Rab11b* | 0.95 | 0.00E+00 | 0.95 | 0.00E+00 | 22.81 | Community_4 |
| *Mreg* | -0.95 | 0.00E+00 | -0.94 | 0.00E+00 | 33.59 | Community_5 |
| *Tiam1* | -0.96 | 0.00E+00 | -0.95 | 0.00E+00 | 31.22 | Community_5 |

| | | | | | | |
|---|---|---|---|---|---|---|
| *Gm11734* | 0.95 | 0.00E+00 | 0.95 | 0.00E+00 | 31.01 | Community_5 |
| *Oplah* | -0.95 | 0.00E+00 | -0.94 | 0.00E+00 | 29.1 | Community_5 |
| *Sucla2* | 0.93 | 0.00E+00 | 0.95 | 0.00E+00 | 24.71 | Community_5 |
| *Nr1d2* | -0.97 | 0.00E+00 | -0.98 | 0.00E+00 | 46.74 | Community_6 |
| *Gm40841* | 0.96 | 0.00E+00 | 0.96 | 0.00E+00 | 32.31 | Community_6 |
| *Tcap* | -0.93 | 0.00E+00 | -0.94 | 0.00E+00 | 28.34 | Community_6 |
| *Per3* | -0.93 | 0.00E+00 | -0.95 | 0.00E+00 | 27.84 | Community_6 |
| *Igf2bp2* | 0.94 | 0.00E+00 | 0.95 | 0.00E+00 | 25.48 | Community_6 |
| *Atp8a1* | -0.98 | 0.00E+00 | -0.97 | 0.00E+00 | 37.83 | Community_7 |
| *Vps13a* | -0.95 | 0.00E+00 | -0.95 | 0.00E+00 | 31.31 | Community_7 |
| *Mylk4* | -0.94 | 0.00E+00 | -0.95 | 0.00E+00 | 27.38 | Community_7 |
| *Cpne2* | 0.94 | 0.00E+00 | 0.93 | 0.00E+00 | 25.84 | Community_7 |
| *Cib2* | -0.96 | 0.00E+00 | -0.94 | 0.00E+00 | 24.22 | Community_7 |

**Table S2: Circadian time-series skeletal muscle datasets analyzed in this study.**

| GEO ID | No. of samples (Muscle) | Organism | Reference |
|---|---|---|---|
| GSE197726 | 54 | *Mus musculus* | (Smith et al., 2023) |
| GSE194106 | 56 | *Mus musculus* | (Xin et al., 2023) |
| GSE273878 | 12 | *Mus musculus* | (Ducharme et al., 2025) |
| GSE308276 | 36 | *Mus musculus* | (Sica et al., 2026) |
| GSE107787 | 36 | *Mus musculus* | (Kinouchi et al., 2018) |
| GSE195724 | 352 | *Mus musculus* | (Kumar et al., 2024) |
| GSE182117 | 186 | *Homo sapiens* | (Gabriel et al., 2021) |
| GSE109825 | 100 | *Homo sapiens* | (Perrin et al., 2018) |
| GSE108539 | 57 | *Homo sapiens* | (Perrin et al., 2018) |
